# The Sequence of 1504 Mutants in the Model Rice Variety Kitaake Facilitates Rapid Functional Genomic Studies

**DOI:** 10.1101/111237

**Authors:** Guotian Li, Rashmi Jain, Mawsheng Chern, Nikki T. Pham, Joel A. Martin, Tong Wei, Wendy S. Schackwitz, Anna M. Lipzen, Phat Q. Duong, Kyle C. Jones, Liangrong Jiang, Deling Ruan, Diane Bauer, Yi Peng, Kerrie W. Barry, Jeremy Schmutz, Pamela C. Ronald

**Author notes:** Address correspondence to and. Corresponding authors: Prof. Pamela C. Ronald, Department of Plant Pathology, University of California, Davis One Shields Avenue, Davis, CA 95616, USA; Dr. Mawsheng Chern, Department of Plant Pathology, University of California, Davis One Shields Avenue, Davis, CA 95616, USA. These authors contributed equally to this work (Guotian Li, Rashmi Jain, Mawsheng Chern).

## Abstract

The availability of a whole-genome sequenced mutant population and the cataloging of mutations of each line at a single-nucleotide resolution facilitates functional genomic analysis. To this end, we generated and sequenced a fast-neutron-induced mutant population in the model rice cultivar Kitaake (*Oryza sativa* L. ssp. *japonica*), which completes its life cycle in 9 weeks. We sequenced 1,504 mutant lines at 45-fold coverage and identified 91,513 mutations affecting 32,307 genes, 58% of all rice genes. We detected an average of 61 mutations per line. Mutation types include single base substitutions, deletions, insertions, inversions, translocations, and tandem duplications. We observed a high proportion of loss-of-function mutations. Using this mutant population, we identified an inversion affecting a single gene as the causative mutation for the short-grain phenotype in one mutant line with a small segregating population. This result reveals the usefulness of the resource for efficient identification of genes conferring specific phenotypes. To facilitate public access to this genetic resource, we established an open access database called KitBase that provides access to sequence data and seed stocks, enabling rapid functional genomic studies of rice.

**One-sentence summary:** We have sequenced 1,504 mutant lines generated in the short life cycle rice variety Kitaake (9 weeks) and established a publicly available database, enabling rapid functional genomic studies of rice.

## INTRODUCTION

Rice (*Oryza sativa*) provides food for more than half of the world’s population, making it the most important staple crop (Gross and Zhao, 2014). In addition to its critical role in global food security, rice also serves as a model for studies of monocotyledonous species including important cereals and bioenergy crops (Izawa and Shimamoto, 1996). For decades, map-based cloning has been the main strategy for isolating genes conferring agronomically important traits (Peters et al., 2003). In Arabidopsis and other model plant species (Alonso et al., 2003; Cheng et al., 2014; Li et al., 2016c), indexed mutant collections constitute highly valuable genetic resources for functional genomic studies. In rice, multiple mutant collections have been established in diverse genetic backgrounds including Nipponbare, Dong Jin, Zhonghua 11, and Hwayoung (Wang et al., 2013b; Wei et al., 2013). Rice mutants have been generated through T-DNA insertion (Jeon et al., 2000; Chen et al., 2003; Sallaud et al., 2003; Wu et al., 2003; Hsing et al., 2007), transposon/retrotransposon insertion (Miyao et al., 2003; Kolesnik et al., 2004; van Enckevort et al., 2005;Wang et al., 2013b), RNAi (Wang et al., 2013a), TALEN-based gene editing (Moscou and Bogdanove, 2009; Li et al., 2012), CRISPR/Cas9 genome editing (Jiang et al., 2013; Miao et al., 2013; Xie et al., 2015), chemical induction, such as ethyl methanesulfonate (EMS) (Henry et al., 2014), and irradiation (Wang et al., 2013b; Wei et al., 2013). Several databases have been established to facilitate use of the mutant collections (Droc, 2006; Zhang, 2006; Wang et al., 2013b). These approaches have advanced the characterization of approximately 2,000 genes (Yamamoto et al., 2012). The usefulness of these rice mutant collections has been hindered by the long life cycle of the genetic backgrounds used (i.e. 6 months) and the lack of sequence information for most of the mutant lines. To address these challenges, we recently established a fast-neutron (FN) mutagenized population in Kitaake, a model rice variety with a short life cycle (9 weeks) (Li et al., 2016b). Here we report the sequence of 1,504 individual lines. We anticipate that the availability of this mutant population will significantly accelerate rice genetic research.

FN irradiation induces a diversity of mutations that differ in size and copy number, including single base substitutions (SBSs), deletions, insertions, inversions, translocations, and duplications (Belfield et al., 2012; Bolon et al., 2014; Li et al., 2016b), in contrast to other mutagenesis approaches that mostly generate one type of mutation (Thompson et al., 2013; Wang et al., 2013b). It generates a broad spectrum of mutant alleles, including loss-of-function, partial loss-of-function and gain-of-function alleles that constitute an allelic series, highly desirable for functional genomic studies. In addition, FN irradiation induces subtle variations, such as SBSs and in-frame insertions/deletions (Indels), which facilitate the study of protein structure and domain functions (Li et al., 2016b). Finally, FN irradiation induces abundant mutations in noncoding genomic regions that may contain important functional transcription units such as microRNAs (Lan et al., 2012) and long noncoding RNAs (Ding et al., 2012). The availability of a FN-induced mutant population with these unique characteristics greatly expands the mutation spectrum relative to other collections and provides researchers the opportunity to discover novel genes and functional elements controlling diverse biological pathways.

Whole-genome sequencing (WGS) of a mutant population, and pinpointing each mutation at a single-nucleotide resolution using next-generation sequencing technologies is an efficient and cost-effective approach to characterize variants in a mutant collection, in contrast to targeting induced local lesions in genomes (TILLING) collections, for which researchers must scan amplicons from a large set of mutants for each use (McCallum et al., 2000). Another commonly used approach to characterize a genome is whole-exome sequencing (WES) (Krasileva et al., 2017). Though it is relatively low-cost, WES does not cover most noncoding regions that potentially contain important functional elements such as microRNAs. Furthermore, WES is unable to identify balanced variants, including inversions and translocations, which are commonly induced by FN irradiation (Biesecker et al., 2011; Li et al., 2016b). Finally, WGS gives more accurate and complete genome-wide variant information than WES, even for the exome (Belkadi et al., 2015). Fully sequenced mutant collections are particularly useful for crops, which have inefficient transformation, and require more time and space for genetic analyses compared to model organisms (Barampuram and Zhang, 2011). Among major crops, rice has the smallest genome (~389 Mb) (Michael and Jackson, 2013), making it the most amenable to WGS, especially with the low cost afforded by sample multiplexing.

In this study, taking advantage of the established FN mutant collection in Kitaake (Li et al., 2016b), we whole-genome sequenced 1,504 lines, identified 91,513 mutations affecting 32,307 genes (58% of all genes in the rice genome) and established the first WGS mutant collection in rice. To facilitate the use of this mutant collection, we established an open access resource called KitBase, which integrates multiple bioinformatics tools and enables users to search the mutant collection, visualize mutations, download genome sequences for functional analysis and order seed stocks.

## RESULTS

### Genome Sequencing

We sequenced 1,504 mutagenized lines, including 1,408 M_2_ lines and 96 M_3_ lines using the Illumina high-throughput sequencing technology, and characterized mutations in these lines. To facilitate downstream analysis, genomic DNA was isolated from a single plant of each line. High-throughput sequencing was performed using the Illumina Hiseq 2000 system, and the resultant sequence reads were mapped to the Nipponbare reference genome using BurrowsWheeler Aligner-Maximal Exact Match algorithm (BWA-MEM) (Li, 2013). On average, 183 million paired-end reads (18.6 Gb) were obtained for each line (Table 1 and Supplemental Data Set 1), and 170 million high-quality reads (93% of the raw reads) were mapped onto the reference genome, giving an average sequencing depth of 45.3-fold for each line. The high sequencing depth of these rice mutant lines facilitated detection of different types of variants.

**Table 1.** Genome Sequencing Summary of Mutagenized Rice Plants Used in This Study

| Summary | Information |
| --- | --- |
| Total samples | 1,504 |
| Mean raw bases (Gb) | 18.6 |
| Mean aligned bases (Gb) | 17.3 |
| Mean sequencing depth (fold)<sup>a</sup> | 45.3 |

<sup>a</sup> The reference genome size of 374 Mb was used in calculating sequencing depth.

### Genomic Variants Detected in the 1,504 Mutant Lines

We used an established variant-calling pipeline containing multiple complementary programs to call variants in each rice line, filtering out variants present in the parental line and those found in two or more rice lines (see Methods). A total of 91,513 FN-induced mutations were detected in the 1,504 rice lines, including 43,483 single base substitutions (SBSs), 31,909 deletions, 7,929 insertions, 3,691 inversions, 4,436 translocations, and 65 tandem duplications (Figure 1 and Supplemental Data Set 2). The largest inversion is 36.8 Mb, the largest tandem duplication 4.2 Mb, and the largest deletion 1.7 Mb (Supplemental Figure 1). To assay the false positive rate, we randomly selected 10 lines and examined all of their mutations (Supplemental Data Set 3). Out of 638 mutation events, we identified 30 false positives (4.7%), indicating that our variant-calling pipeline is robust. 60% of these false positives are either SBSs or small Indels (<30bp), mostly in the polynucleotide or repetitive regions. Only 4 false positives out of 638 mutations events (0.6%) are in coding regions, indicating the minimal impact of false positives on mutated genes.

**Figure 1.**
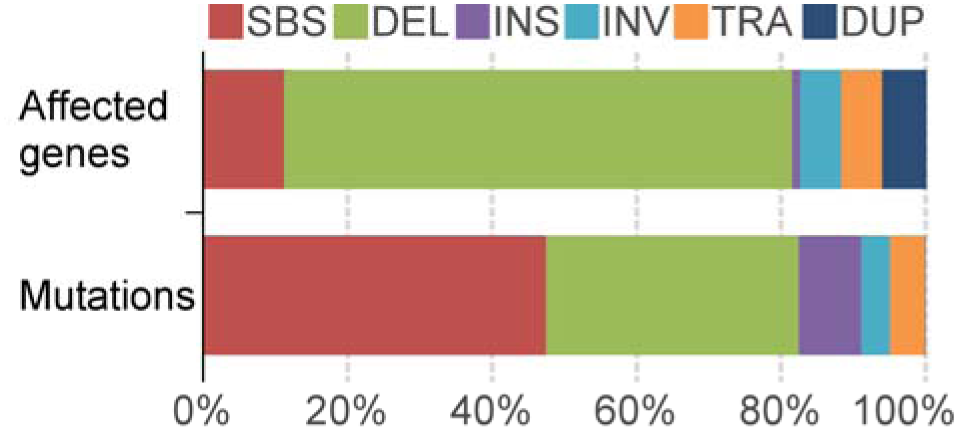
Mutations and Affected Genes in the Kitaake Rice Mutant Population. SBS, single base substitutions; DEL, deletions; INS, insertions; INV, inversions; TRA, translocations; and DUP, tandem duplications.

Among the 91,513 mutations, SBSs are the most abundant variants, accounting for 48% of mutation events. We identified 48,030 non-SBS mutations, of which deletions account for 66%. Small deletions make up the majority of all deletion events: deletions smaller than 100 bp account for nearly 90% of all deletions (Table 2). There are 7,469 single base deletions, accounting for 23% of all deletion events. The average deletion size is 8.8 kb.

**Table 2.** Size Distribution of Deletions in the Kitaake Rice Mutant Population

| Size | Number | Average size | Percentage |
| --- | --- | --- | --- |
| 1-10 bp | 21,998 | 3.7 bp | 68.9 |
| 10-100 bp | 6,588 | 21.7 bp | 20.6 |
| 100 bp-10 kb | 1,274 | 2.5 kb | 4.0 |
| 10 kb-1 Mb | 2,029 | 124.3 kb | 6.4 |
| > 1 Mb | 20 | 1.2 Mb | 0.1 |
| Total | 31,909 | 8.8 kb | 100.0 |

To analyze the distribution of mutations in the genome, all mutations from the sequenced lines were mapped to the reference genome (Figure 2). We found that the FN-induced mutations are distributed evenly across the genome, except for some repetitive regions with low mapping quality reads or no read coverage caused by the inability to confidently align the reads to the reference. Many translocations were identified in the mutant population, shown by the connecting lines (Figure 2E). The density of translocations is similar on each chromosome, ranging from 20.4/Mb to 26.8/Mb (Supplemental Table 1). The genome-wide mutation rate of the Kitaake rice mutant population is 245 mutations/Mb. The even distribution of FN-induced mutations is similar to the distribution of mutations generated through chemical mutagenesis of sorghum and *Caenorhabditis elegans* (Thompson et al., 2013; Jiao et al., 2016).

**Figure 2.**
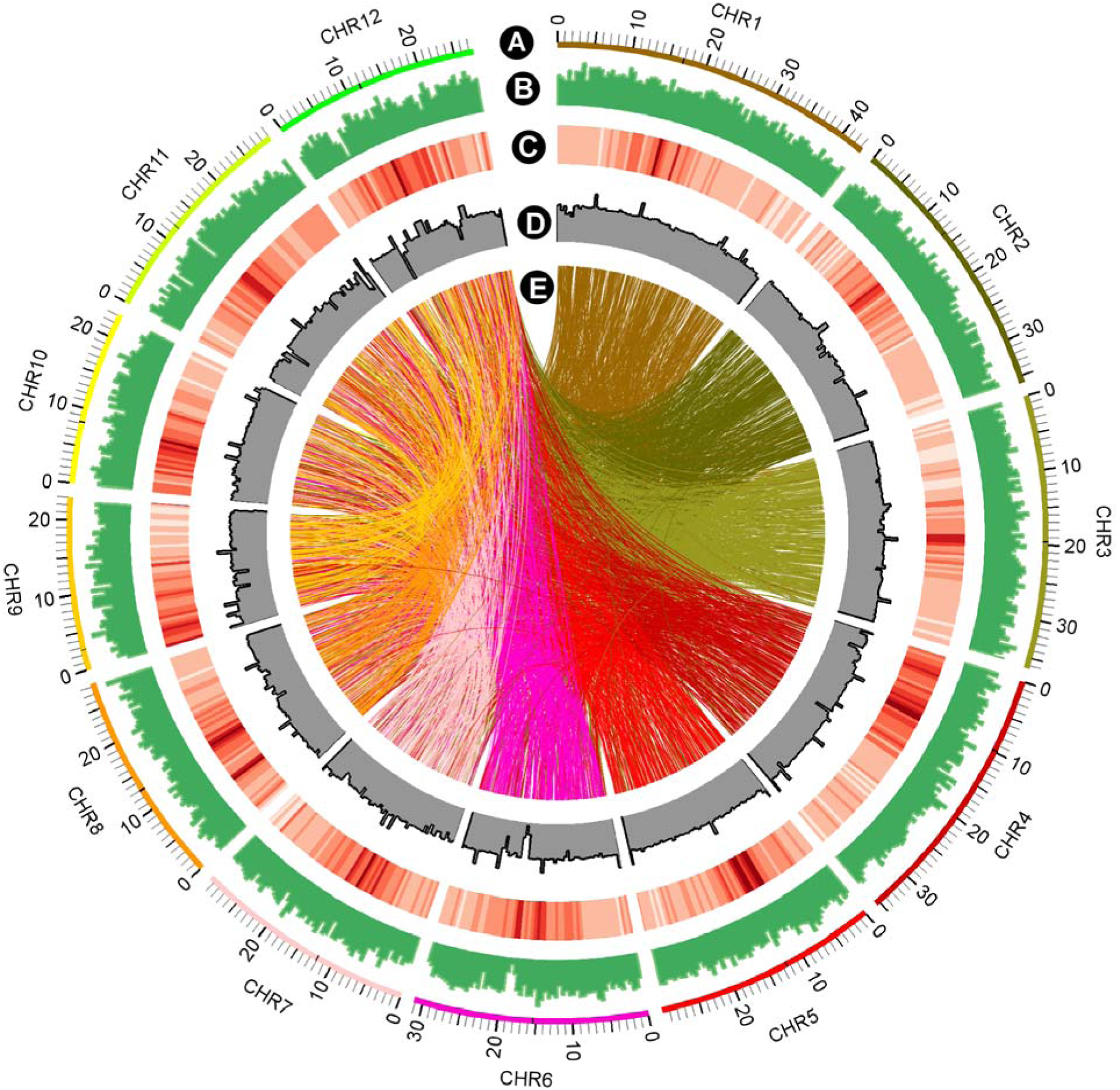
Genome-Wide Distribution of FN-Induced Mutations in the Kitaake Rice Mutant Population. **(A)** The twelve rice chromosomes represented on an Mb scale. **(B)** Genome-wide distribution of FN-induced mutations in non-overlapping 500 kb windows. The highest column equates to 242 mutations/500kb. **(C)** Repetitive sequences in the reference genome in non-overlapping 500 kb windows. The darker the color, the higher perentage content of repetitive sequences. **(D)** The sequencing depth of the parental line X.Kitaake. The highest column indicates 300 fold. **(E)** Translocations. Translocations are represented with connecting lines in the color of the smaller-numbered chromosome involved in the translocation.

### Genes Affected in 1,504 Mutant Lines

Genes affected by FN-induced mutation were identified using an established pipeline (see Methods). A total of 32,307 genes, 58% of all 55,986 rice genes (Kawahara et al., 2013) are affected by different types of mutations (Figure 1 and Supplemental Data Set 4). Deletions affect the greatest number of genes, 27,614, accounting for 70% of the total number of affected genes. SBSs, constituting the most abundant mutation, only affect 4,378 genes (11%). Inversions, translocations, and duplications affect 2,230, 2,218, and 2,378 genes, respectively.

To test whether the affected genes are biased with respect to a particular biological process, we used gene ontology (GO) analysis to classify all affected genes into major functional categories (Ashburner et al., 2000; Du et al., 2010). As expected, the selected biological process categories “DNA metabolic process”, “protein modification process”, and “transcription” have the most hits and show similar percentages to the mutation saturation (58%) (Supplemental Table 2 and Supplemental Figure 2). We observed that the terms of “DNA metabolic process” and “cellular component organization” show slightly higher percentages within the biological process category, whereas “photosynthesis”, and “transcription” show much lower percentages (Supplemental Table 2). Core eukaryotic genes are highly conserved and are recalcitrant to modifications (Parra et al., 2008). We analyzed a set of core eukaryotic genes and showed that 40% of these analyzed are affected, mostly by heterozygous mutations (Supplemental Data Set 5). Taken together, these results suggest that, although FN-induced are evenly distributed across the genome in the mutant population, the affected genes are biased against mutations in core gene functions.

### FN-Induced Mutations in Each Rice Line

To assess the overall effect of FN irradiation in each sequenced line, the mutations and genes affected in each line were calculated (Supplemental Data Set 1). On average, each line contains 61 mutations. The distribution of the number of mutations per line corresponds to a normal distribution (Figure 3). Of the 1,504 lines, 90% have fewer than 83 mutations per line (Figure 3). The average number of genes affected per line is 43 (Supplemental Data Set 1). The variation of affected genes per line is greater than that of mutations per line (Table 3), due to the presence of large mutation events (Supplemental Data Set 4). For example, line FN-259 has the most genes affected (681 genes) in this mutant population, largely due to the 4.2 Mb tandem duplication that affects 667 genes (Supplemental Data Set 4). However, 76% of the mutated lines contain no more than 50 mutated genes per line (Table 3). Only 10% of the mutated lines contain more than 100 affected genes. The relatively low number of mutations per line for most lines in the Kitaake rice mutant population facilitates downstream cosegregation assays.

**Figure 3.**
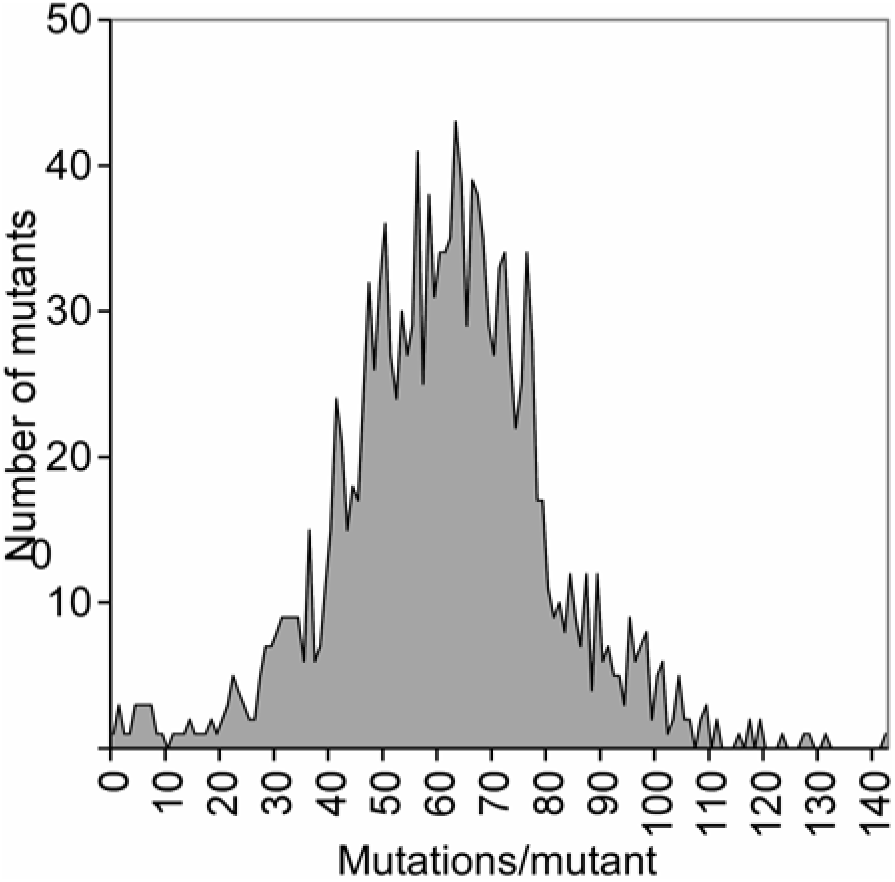
Distribution of the Number of Mutations per Line in the Kitaake Rice Mutant Population. The x-axis represents the number of mutations per line. The y-axis indicates the number of mutants containing the indicated number of mutations.

**Table 3.** Affected genes per Line in the Kitaake Rice Mutant Population

| Effect Type | Genes | Percentage |
| --- | --- | --- |
| Start lost | 7 | 0.0 |
| Splice site | 52 | 0.6 |
| Stop gained/lost | 303 | 3.4 |
| Frameshift<sup>a</sup> | 4,103 | 46.6 |
| Truncation<sup>b</sup> | 4,348 | 49.3 |
| Total<sup>c</sup> | 8,221 | 100.0 |

### Loss-of-Function Mutations

A large number of loss-of-function mutations were identified in this mutant population. Loss-of-function mutations completely disrupt genes. They are of considerable value in functional genomics because they often clearly indicate the function of a gene (MacArthur et al., 2012). To identify loss-of-function mutations from the Kitaake rice mutant population, we adopted the definition as described (MacArthur et al., 2012) with minor modifications: we included mutations affecting start/stop codons and intron splice sites as well as mutations causing frameshifts, gene knockouts or truncations (See Methods). There are 28,860 genes affected by loss-of-function mutations (Figure 4 and Supplemental Data Set 6), accounting for 89% of the genes affected in this mutant population and 52% of all rice genes in the genome. The 344 genes affected by loss-of-function SBSs account for 1% of all genes mutated by all loss-of-function mutations. In contrast, loss-of-function deletions disrupt 26,822 genes, accounting for 84% of genes mutated by loss-of-function mutations. Inversions and translocations disrupt 2,230 and 2,218 genes, respectively. These results explicitly show that FN irradiation induces a high percentage of loss-of-function mutations and that deletions are the main cause.

**Figure 4.**
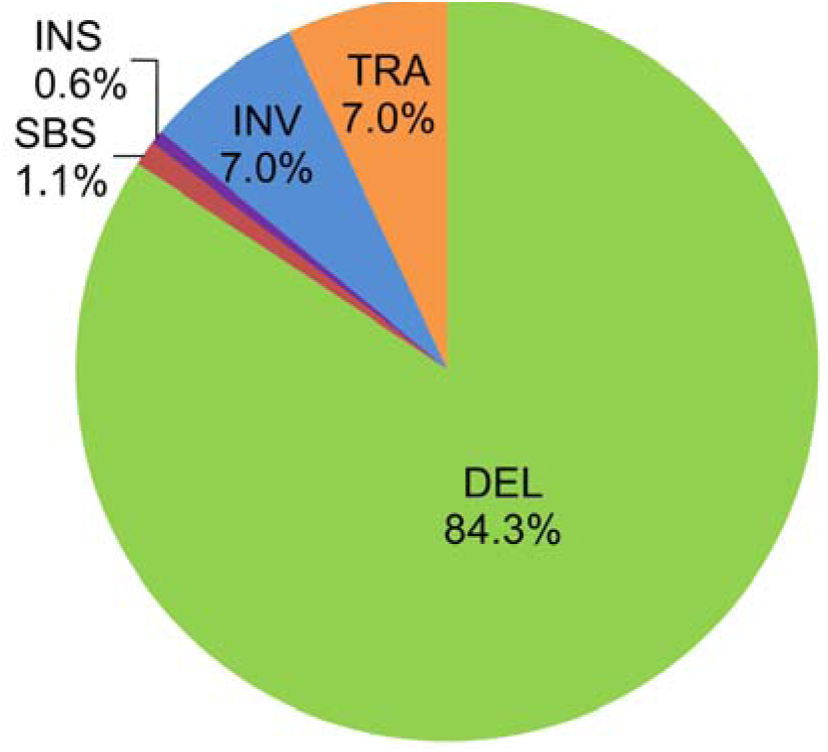
Genes Mutated by Loss-of-Function Mutations in the Kitaake Rice Mutant Population. The percentage of gene mutated by each type of mutation is shown. DEL, deletions; TRA, translocations; INV, inversions; INS, insertions; and SBS, single base substitutions. Genes affected by tandem duplications, the copy number of which is increased, are not included.

Loss-of-function mutations affecting a single gene allow straightforward functional genomic analysis. We analyzed genes affected by these mutations and cataloged them according to the effect of the mutation, and identified 8,221 such genes (Table 4 and Supplemental Data Set 7). Frameshifts and truncations, mostly a result of deletions, inversions and translocations, account for 96% of the genes, which indicates the importance of these non-SBS variants.

**Table 4.** Genes Mutated by Loss-of-Function Mutations Affecting a Single Gene

| Genes/mutant | Mutants | Percentage |
| --- | --- | --- |
| <50 | 1,142 | 76 |
| 50-100 | 215 | 14 |
| >100 | 147 | 10 |
| Total | 1,504 | 100 |

<sup>a</sup> A frameshift refers to Indels, although it has a truncation effect on the gene.
<sup>b</sup> The breakpoint of the loss-of-function mutation falls in the genic region or the gene is completely deleted due to structural variants.
<sup>c</sup> Only includes unique genes. This number is smaller than the sum of genes affected in each category as one gene can be affected by different types of mutations.

### FN-Induced Single Base Substitutions

To draw comparisons between the FN-induced and EMS-induced mutant populations, we conducted a detailed analysis of SBSs. There is an average of 29 SBSs per line (Supplemental Figure 3). Ninety percent of our lines contain between 10 and 50 SBSs per line. There are 118 SBSs in mutant FN1423-S, the highest number of SBSs per line in the mutant population. SBSs are evenly distributed in the genome (Supplemental Figure 4), similar to the EMS-induced mutant populations (Thompson et al., 2013; Jiao et al., 2016). 37.9% of SBSs map within genes and 62.1% to intergenic regions (Supplemental Table 3). Of the genic SBSs, 17.3% are within exons, 17.4% within introns, 3.2% within untranslated regions (UTRs), and 0.1% at canonical splice sites (GT/AG). Non-synonymous SBSs, which represent 12.4% of all SBSs, are found in 4,378 genes (Supplemental Data Set 4). Of these, 11.5% cause missense mutations, 0.8% cause nonsense mutations, and 0.1% result in readthrough mutations (Supplemental Table 3).

The amino acid changes of the three mutant populations were further analyzed using heat maps (Figure 5A). The amino acid changes of the FN-induced Kitaake rice mutant population are relatively evenly distributed, compared to the two EMS-induced mutant populations (Figure 5B, C). The differences are due to the less biased nucleotide changes of the FN-induced mutant population compared to the two EMS-induced mutant populations (Figure 5D). The frequency of the most common GT>AC nucleotide changes in the FN-induced mutant population is 42.5%, half that in the EMS-induced population (88.3%) (Henry, 2014) (Figure 5D). All possible amino acid changes caused by a single nucleotide change are present in the FN-induced mutant population (Figure 5A). Alanine to threonine or valine changes show a much higher frequency, 4.5% and 4.3%, respectively, compared to the average amino acid change frequency of 0.7%. Alanine to threonine or valine changes occur so often because these three amino acids are all encoded by four codons, and a single nucleotide change (GT>AC), the most common nucleotide changes in the mutant population, is enough to change the amino acid (Figure 5E). Similar patterns are found in the two EMS-induced mutant populations (Thompson et al., 2013; Jiao et al., 2016). Some amino acid changes occur infrequently, because the occurrence frequency of these amino acids is low in rice (Itoh et al., 2007) and/or a single GT>AC change may not be sufficient to cause the amino acid change. The results demonstrate that FN irradiation induces diverse amino acid changes at higher frequencies than EMS treatment and that FN irradiation can result in amino acid mutations rarely achieved by chemical mutagens.

**Figure 5.**
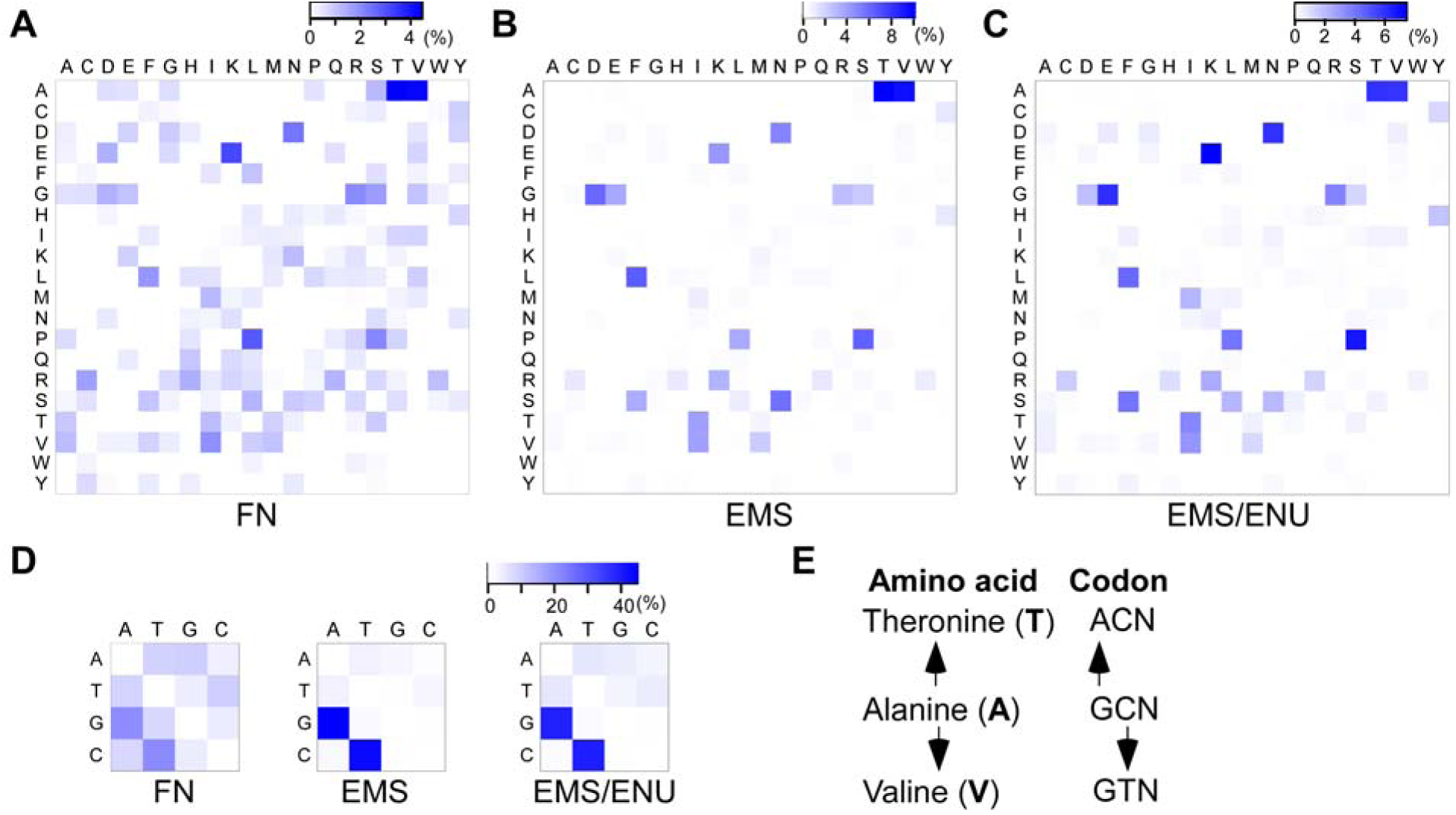
Amino Acid and Nucleotide Changes in the FN- and Two EMS-Induced Mutant Populations. **(A)** Amino acid changes in the FN-induced Kitaake rice mutant population. The single letter symbol of amino acids is labeled in heat maps **(A), (B)** and **(C).** Each cell is colored according the percentage of the specific amino acid change compared to all the amino acid changes in the mutant population. The blank cells in **(A)** represent amino acid changes that require alterations of two or three nucleotides in the codon. **(B)** Amino acid changes in the ethyl methanesulfonate (EMS)-induced mutant population in the rice Nipponbare (Henry et al., 2014). **(C)** Amino acid changes in the EMS/N-ethyl-N-nitrosourea (ENU)-induced mutant population in *C. elegans.* This population was generated with either EMS, ENU, or a combination of both (Thompson et al., 2013). **(D)** Nucleotide changes in the FN-induced Kitaake rice mutant population (left), the EMS-induced mutant population in the rice Nipponbare (middle), and the EMS/ENU-induced mutant population in *C. elegans* (right). Nucleotides are labeled in heat maps. Each cell is colored according the percentage of the specific nucleotide change compared to all the nucleotide changes in the mutant population. Only nucleotide changes that cause missense mutations are included. **(E)** The most frequent amino acid changes in the three induced mutant populations. The codon changes show that nucleotide changes of alanine (A) to threonine (T) or to valine (V) are in the conserved GC>AT changes. Single letters of amino acids are shown in bold, and nucleotides are not. N stands for nucleotides A, T, C, and G.

### An Inversion in Mutant FN1535 Cosegregates with the Short Grain Phenotype

Grain shape is a key determinant of rice yield (Huang et al., 2013). When growing the mutated lines, we observed that line FN1535 produces significantly shorter grains compared to the parental line (Figure 6). The mutant is also dwarfed and shows a much shorter panicle. In a segregating population, we observed 34 normal plants and 13 short-grain plants, a 3:1 ratio. A goodness-of-fit test based on χ^2^ analysis of the phenotypic ratio revealed that the observed values are statistically similar to the expected values, indicating that the short-grain phenotype is likely caused by a recessive mutation. Next, we identified all mutations in line FN1535. We identified 76 mutations, including 26 SBSs, 38 deletions, 10 insertions, and 2 inversions (Supplemental Data Set 2). These mutations affect seven non-transposable element (TE) genes (Supplemental Table 4). To identify which mutation is responsible for the short-grain phenotype, we prioritized them based on their putative loss-of-function effects and predicted functions of the affected genes. We prioritized a 37 kb deletion on chromosome 7 that affects 5 genes, an inversion on chromosome 5 affecting one gene, and a SBS on chromosome 6 that affects one gene. Using the segregating population of 50 plants, we found that the inversion on chromosome 5, not the chromosome 7 deletion or the chromosome 6 SBS, cosegregates with the phenotype (Figure 6D and Supplemental Figure 6). We analyzed the causative inversion in detail. One breakpoint of the inversion is in the fourth exon of gene LOC_Os05g26890, which truncates the gene (Figure 6E). The other breakpoint of the inversion is not in the genic region. This gene, named *Dwarf 1/RGA1*, was previously isolated using a map-based cloning strategy (Ashikari et al., 1999). Gene *Dwarf 1/RGA1* encodes a Gα protein, which is involved in gibberellin signal transduction (Ueguchi-Tanaka et al., 2000). Mutations in gene *Dwarf 1/RGA1* cause the dwarf and short-grain phenotypes (Ashikari et al., 1999). Identical phenotypes were observed in line FN1535 (Figure 6). These results demonstrate that we can rapidly pinpoint the genetic lesion and gene conferring a specific phenotype using a small segregating population of the mutant line.

**Figure 6.**
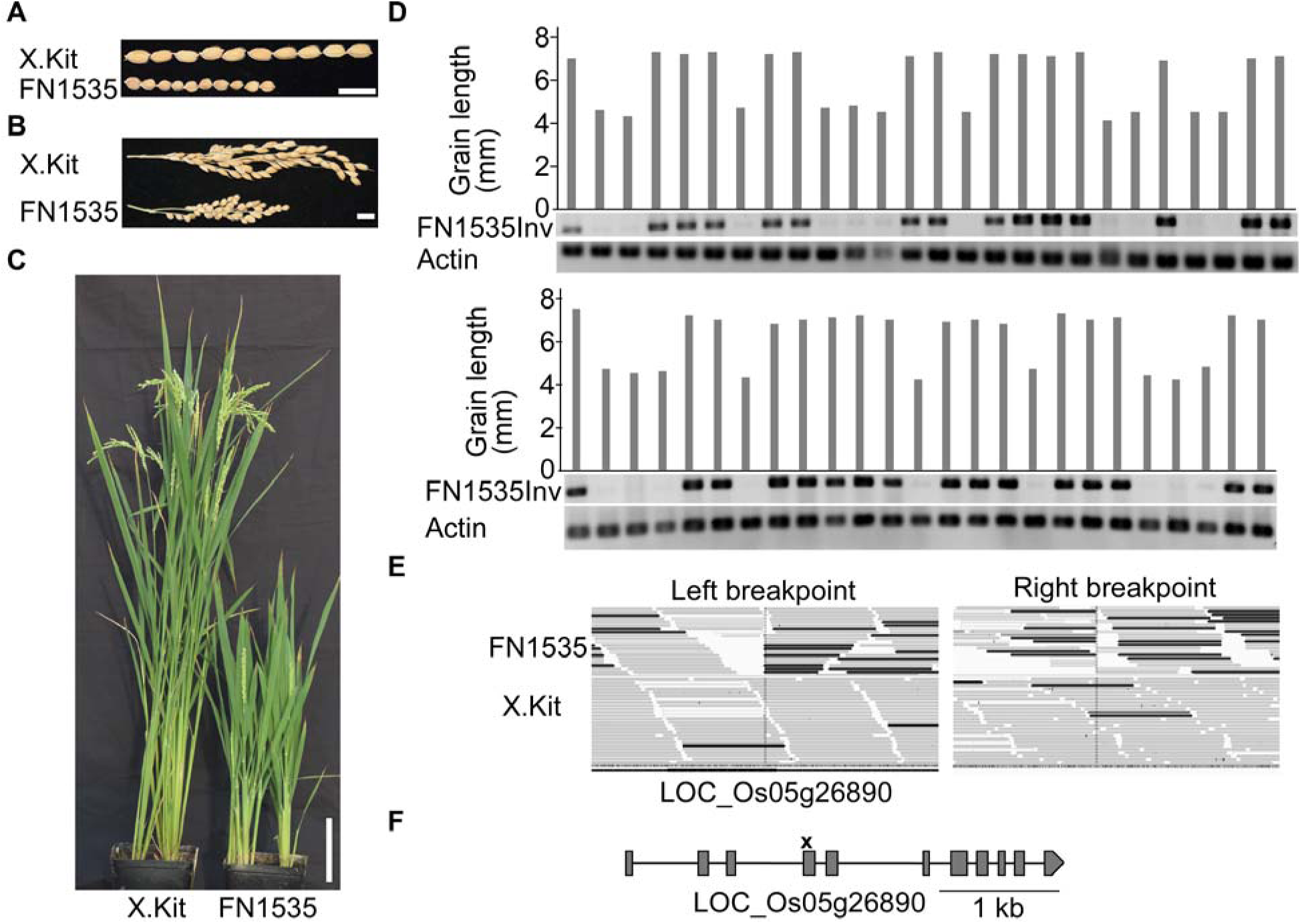
An Inversion Cosegregates with the Short-Grain Phenotype in Line FN1535. **(A)** Seeds of line FN1535 and the nonirradiated parental line X.Kitaake (X.Kit). Bar = 1 cm. **(B)** Panicles of line FN1535 and the parental line X.Kit. Bar = 1 cm. **(C)** Line FN1535 and the parental line X.Kit at the grain filling stage. Bar = 10 cm. **(D)** The inversion on chromosome 5 of line FN1535 cosegregates with the short-grain phenotype. Grain length was measured by lining up 10 mature seeds of each plant as shown in (A), and the average grain length was calculated. The first lane of the top panel represents the parental line X.Kit. Fifty progeny used in the cosegregation analysis were represented in two panels. FN1535Inv indicates the PCR results targeting the inversion on chromosome 5 of line FN1535. A band indicates the presence of at least one parental allele in the plant. Actin primers were used for the DNA quality control. **(E)** Integrative Genomics Viewer (1GV) screenshots of the two breakpoints of the inversion on chromosome 5 of line FN1535. The dark color indicates the anomalous reads of the inversion. Only the left breakpoint affects a gene (LOC_Os05g26890). X.Kit indicates the parental line. **(F)** Gene structure of LOC_Os05g26890. The breakpoint of the inversion is marked with a cross symbol. Gray boxes indicate exons, and lines for introns. The gene structure diagram is modified from the Nipponbare reference genome.

### Access to Mutations, Sequence Data and Seed Stocks

Publicly available access to high-throughput resources are essential for advancing science (McCouch et al., 2016). To make the mutant collection and associated data available to users, we established an open access web resource named KitBase (http://kitbase.ucdavis.edu/) (Figure 7). KitBase provides the mutant collection information, including sequence data, mutation data, and seed information for each rice line. Users can use different inputs, including gene IDs, mutant IDs, and DNA or protein sequences to search and browse KitBase (Figure 7A). Search with DNA or protein sequences will be carried out with the standalone BLAST tool (Deng et al., 2007). Both MSU LOC gene IDs and RAP-DB gene IDs (Kawahara et al., 2013; Sakai et al., 2013) can be used in searching the database. Mutations are visualized using the web-based interactive JBrowse genome browser, in which different symbols are used to indicate different types of mutations at the corresponding locations. Users interested in a particular region of the genome can browse all the mutations from KitBase in that region (Figure 7B). This visual approach enables users to identify multiple allelic mutations and elucidate gene function quickly. Mutation information for each line can be downloaded from KitBase. The original sequence data and primary mutation data of lines in KitBase can be accessed through the National Center for Biotechnology Information (NCBI) and the Joint Genome Institute (JGI) (Supplemental Data Set 1). A seed request webpage was set up for seed distribution with a minimal handling fee. The seed distribution is currently subsidized by the Department of Energy via the Joint BioEnergy Institute. The user-friendly genetic resources and tools in this open access platform will facilitate rice functional genomic studies.

**Figure 7.**
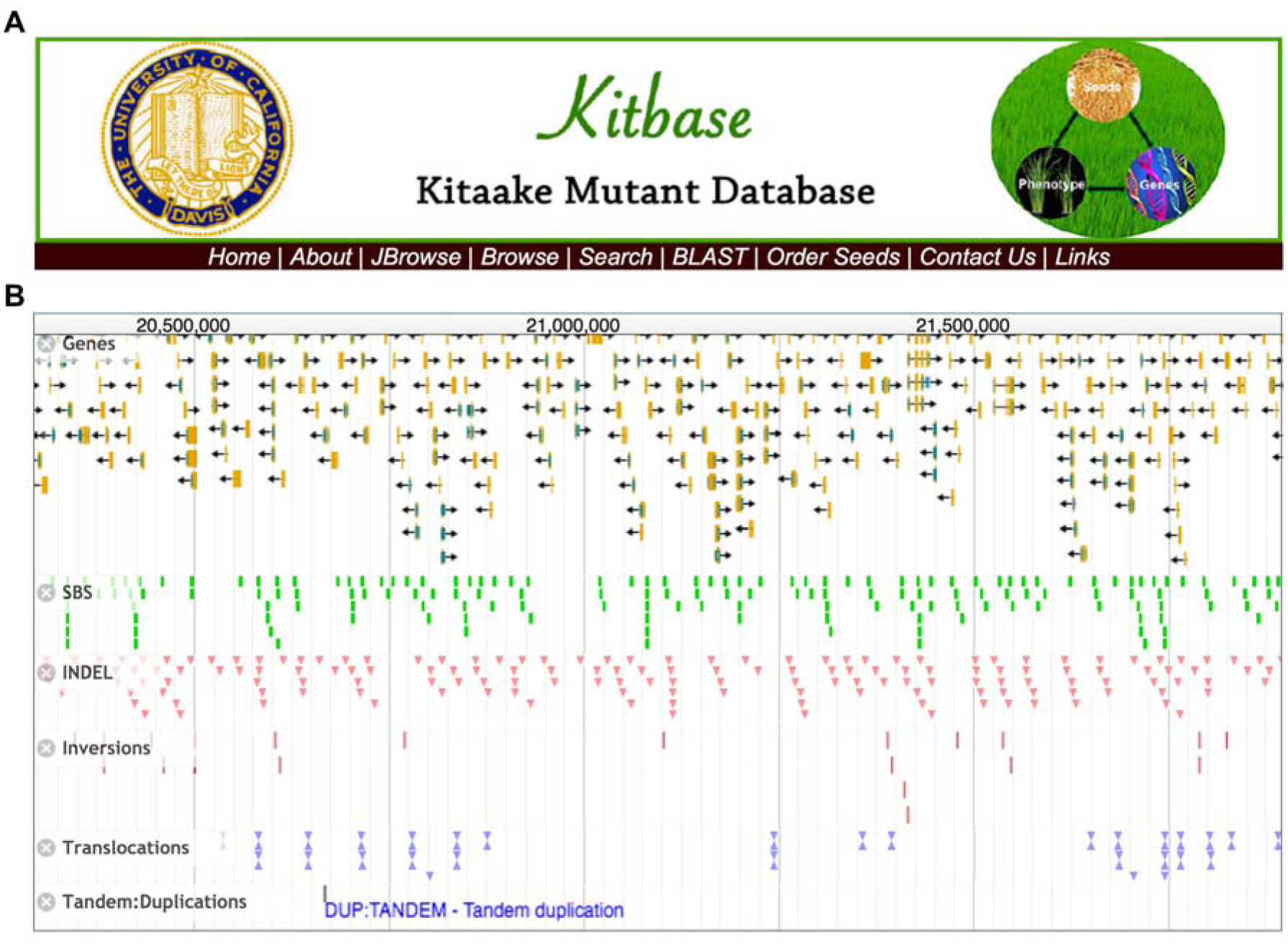
The Navigation Page and Tools in KitBase. **(A)** The main navigation page of KitBase. KitBase can be queried using either mutant ID, MSU7 LOC gene ID, or RAP-DB gene ID. Both DNA and protein sequences can be used as the input in BLAST search. **(B)** A JBrowse snapshot of mutations in a genomic region of the mutant population.

## DISCUSSION

We describe a new resource that facilitates functional genomic studies of rice. A key technical feature of our mutant collection is the low level of mutagenesis (Li et al., 2016b). There is an average of 61 mutations per line (Figure 3), which means that only a small segregating population is needed to identify the causative mutation, for example, 50 plants as demonstrated by our study of the short-grain phenotype. Similar approaches have been used in Arabidopsis and other organisms to clone genes from WGS lines with a small population (Schneeberger, 2014; Li et al., 2016a). In contrast, a large segregating population is required to identify the causative mutation using conventional genetic mapping approaches. Our population requires 0-1 round of backcross. In contrast, some heavily mutagenized populations that carry thousands of SBSs in each mutant line require multiple rounds of time-consuming backcrosses to clean up the background of the line (Jiao et al, 2016). Because we sequenced a single plant instead of pooled samples, users can readily identify segregating populations to pinpoint the mutation responsible for the phenotype often without carrying out backcrossing. We estimate that 67% of all mutations in the M_2_ sequenced lines are heterozygous. For these heterozygous mutations, the progeny seeds available in KitBase can be directly used for cosegregation analysis. For homozygous mutations (33% of detected mutations), the sibling plants of the sequenced lines or progeny of their sibling plants that carry the corresponding heterozygous mutations can be used for cosegregation analysis (Figure 7), which significantly expedites genetic analysis. Users can also backcross the mutant to the parental line to create segregating progeny if needed. Compared to other sequence-indexed mutant populations including the T-DNA or Tos17 populations, WGS detects all possible variants, regardless whether the variant is induced or spontaneous, tagged or not, which avoids the problem of somatic variants going undetected even when the tag is clearly identified in some mutant populations (Wang et al., 2013b). The public availability of the mutant population in the early flowering, photoperiod insensitive Kitaake variety will lower the threshold for researchers outside the rice community to examine functions of their gene of interest in rice.

FN irradiation induces a high proportion of loss-of-function mutations, which means that a relatively small population is needed to mutate all the genes in the genome. In 1,504 mutated lines, 89.3% of all the affected genes are mutated by loss-of-function mutations (Figure 4). In comparison, only 0.2% of the EMS-induced mutations are annotated as loss-of-function mutations in the sequenced sorghum population (Jiao et al., 2016). 80,000 T-DNA insertion rice lines are needed to reach the same mutation saturation level (58%), without taking into account that T-DNA insertions are biased to certain genomic regions (Wang et al., 2013b). Many screens can only be performed when plants are mature, such as yield-related traits (Figure 7A); this means a serious delay when a variety with a long life cycle is used. The Kitaake rice mutant population enables researchers to do studies and complete screens on a relatively small population in a much shorter time. These features make it easier for researchers to conduct studies on complex traits like yield and stress tolerance, which were once too time- and labor-intensive. In addition, with FN-induced loss-of-function mutations, researchers also avoid the variation in knockdown efficiency or off-target issues with approaches such as RNAi or CRISPR-Cas9 (Peng et al., 2016).

Structural variants (variants>1 kb) are known to be the cause of some human diseases, such as the well-known Down and Turner syndromes, and are associated with several cancers (Weischenfeldt et al., 2013; Carvalho and Lupski, 2016). Limited studies in plants show that structural variants contribute to important agricultural and biological traits, like plant height, stress responses, crop domestication, speciation, and genome diversity and evolution (Lowry and Willis, 2010; Huang et al., 2012; Saxena et al., 2014; Zmienko et al., 2014; Zhang et al., 2015; Zhang et al., 2016). However, the study of structural variants in plants is still challenging because they are often identified in different plant varieties/accessions, and the numerous variants between varieties/accessions complicate the study of function of a specific structural variant (Saxena et al., 2014; Zhang et al., 2016). Our Kitaake rice mutant population provides structural variants in the same genetic background, with only a few of structural variants per line, significantly facilitating the study of the function and formation of structural variants in plants (Supplemental Data Set 2).

One limitation of this Kitaake rice mutant population is that large deletions cause loss of function of many genes at once. Although such large deletions are important in achieving saturation of the genome and are valuable in screens, they also pose challenges. A large deletion is likely homozygous lethal, and lethality makes it hard to study genes in the large deletion. In addition, if a large deletion is identified as the causative mutation, determining which gene causes the phenotype requires multiple complementation tests (Wei et al., 2013; Chern et al., 2016). However, as more mutagenized rice lines are collected, multiple lines carrying independent mutations of the same gene will allow researchers to quickly identify the gene associated with the phenotype (Henry et al., 2014). Another approach is to search other mutant collections to identify mutations in individual genes and connect the gene with the phenotype. Another deficit of the current mutant population is the lack of enough mutant alleles in core eukaryotic genes and genes involved in “photosynthesis” and “developmental process” (Supplemental Table 2 and Supplemental Data Set 5), which is likely due to the lethality of these genes and the high portion of loss-of-function mutations induced by FN irradiation. Other rice mutant collections, for example, the EMS-induced mutant populations, would be complementary on this aspect by providing alleles with less severe effects on these genes (Krishnan et al., 2009; Henry et al., 2014). Though we have sequenced the rice lines at a high depth (45-fold), it is still challenging to accurately call dispersed duplications that might result from imbalanced translocations; therefore we include only tandem duplications. Owing to the nature of variant calls made by the algorithms we used, the genotype (homozygosity/heterozygosity) of large structural variants is not included. However, users can use tools such as IGV (Robinson et al., 2011) to obtain the genotype information with available mutant files from KitBase (Figure 6). Cost is another factor to consider when using WGS in profiling variants in a population, though this consideration is not specific to the Kitaake mutant population. It still initially requires a considerable investment when establishing a WGS population but the price of sequencing has dropped dramatically with the technological improvement (Goodwin et al., 2016). One approach to alleviate the financial challenge is through community collaboration, as a WGS population greatly benefits every researcher in that community.

A systematically phenotyped WGS mutant population is highly desirable for functional genomic studies and can rapidly bridge the genotype-to-phenotype knowledge gap. The Kitaake rice mutant population we describe in this study paves the way toward the genomics-phenomics approach in functional genomics. The recently developed high-throughput phenotyping platform makes it feasible to conduct large-scale phenotyping in rice (Yang et al., 2014). We anticipate that adding systematic phenotypic data to these WGS lines will significantly boost the utilization of the mutant collection in this model rice variety. Pairing our genomics resource with a high-throughput phenomics platform will greatly expand the capacity of researchers in rice functional genomic studies.

This study provides a cost-efficient and time-saving open access resource to gene discovery in a short life cycle rice variety by integrating physical mutagenesis, WGS, and a publicly available online database. With the WGS approach, crops are advantageous compared to some mammalian systems, because a sufficiently large mutagenized population can be easily generated and maintained as seed stocks at a low cost, and the mutagenized lines can be directly planted and screened on a large scale in the field. Furthermore, as physical mutagenesis is not considered a transgenic approach, mutants with elite traits from the screens can be directly used in breeding. Given the close phylogenetic relations of rice to other grasses (Devos and Gale, 2000), this resource will also facilitate the functional studies of other grasses, such as cereals and candidate bioenergy crops (Yuan et al., 2008).

## METHODS

### Plant Materials and Growth

The mutagenized lines used in this study were generated using fast-neutron (FN) irradiation described previously (Li et al., 2016b). Briefly, 10,000 rice seeds of the parental line X.Kitaake, a line of the *japonica* cv. Kitaake carrying the XA21 gene under control of the maize ubiquitin promoter, were mutagenized at 20 grays of irradiation (Li et al., 2016b). Over 7,300 fertile M_1_ lines constitute the mutant population. The sequenced plants are mainly derived from the M_2_ generation and some from the M_3_ generation (Supplemental Data Set 1). The seeds from each line were dried and stored. To collect leaf tissues for DNA isolation, seeds were soaked in water in petri dishes at 28°C in a growth chamber for one week and then transplanted to an environmentally-controlled greenhouse at the University of California, Davis. In the greenhouse, light intensity across the spectrum from 400 to 700 nm was approximately 250 μmol m^−2^s^−1^ and the temperature was set to 28–30 °C and humidity to 75–85%. During November to April, artificial lights were supplemented to maintain the light intensity and the day/night period to 14/10 (Schwessinger et al., 2015).

### DNA Sequencing and Read Mapping

DNA isolation and sequencing were done as described previously (Li et al., 2016b). Briefly, the young leaf tissue was sampled with liquid nitrogen from a three-week-old plant of each line and then stored in the −80°C freezer for DNA isolation. High-quality genomic DNA was isolated from young leaves using the cetyltrimethyl ammonium bromide (CTAB) method (Xu et al., 2012). DNA was quantified using Nanodrop (Thermo Scientific) and fluorometer (Tecan) with the PicoGreen dsDNA assay kit (Life Technologies). The integrity of DNA samples was assayed by running samples through a 0.7% agarose gel. Only high-quality DNA was used in sequencing. Sequencing was performed on the HiSeq 2000 sequencing system (Illumina) at the Joint Genome Institute (JGI) following the manufacturer’s instructions. Sequencing was targeted to a minimum sequencing depth of 25-fold for each rice line to facilitate the downstream variant detection. The 2×100 bp paired-end sequence reads were mapped to the Nipponbare genome version 7 (Kawahara et al., 2013) using the mapping tool Burrows-Wheeler Aligner-MEM (BWA version 0.7.10) with default parameters (Li, 2013). The 41 mutant lines published in the pilot study were also included (Li et al., 2016b).

### Genomic Variant Detection

Genomic variant detection was conducted as described in (Li et al., 2016b) with minor modifications. Samples were analyzed in groups of no more than 50 mutant lines including the nonirradiated control line, given the high computational requirement of handling such a large data set. Genomic variants were called using a set of complementary tools, including SAMtools (Li and Durbin, 2009), BreakDancer (Chen et al., 2009), Pindel (Ye et al., 2009), CNVnator (Abyzov et al., 2011), and DELLY (Rausch et al., 2012). For the results from each tool, we removed all variants detected in the parental genome and those found in two or more samples in that group. We then merged results from each tool by filtering out redundant records. SAMtools and Pindel were used to call SBSs and small Indels (<30bp). The minimum phred scaled quality score of variants called by SAMtools was set to 100. Pindel version 0.2.4 was run with default parameters using BreakDancer results as the input. Small Indel results detected by Pindel were filtered with three criteria: 1) the variant site had at least 10 reads, 2) at least 30% of the reads supported the variant, and 3) the control line had at least 50 reads as described (Li et al., 2016b). Large variants (≥30bp) were called using BreakDancer, Pindel, CNVnator, and DELLY as described in (Li et al., 2016b). For large variants, Pindel results were filtered using the criteria listed above. Pindel sometimes reports the same common variant at multiple close positions in different samples. Therefore, we merged these events if the distance between the variants was less than 10 bp. We used a bin size of 1 kb for CNVnator to detect large deletions (≥30bp). Inversion and translocation results were used from DELLY. Due to the nature of variant calls made by the algorithms (Ye et al., 2009), our results only included tandem duplications but not dispersed duplications. Only tandem duplications from Pindel were used and further filtered based on read depth variance. The false positive rate was calculated by manually examining all mutations *in silico* using Integrative Genomic Viewer (IGV) (Robinson et al., 2011) from 10 randomly selected samples. Snapshots of mutations were generated using IGV unless stated otherwise. The mutation density was calculated by adding up all mutations from the mutant population in every non-overlapping 500 kb window for each chromosome. The genome-wide distribution of mutations was drawn using Circos version 0.66 (Krzywinski et al., 2009).

### Functional Annotation of Mutations

SnpEff (Yang et al., 2015) was used to annotate functional effects of the mutation based on the reference genome version 7 (Kawahara et al., 2013). Genes affected by each type of mutation were further analyzed using specific approaches as described (Li et al., 2016b). Briefly, we only include missense mutations and SBSs affecting the start/stop codon or the canonical GT/AG intron splicing sites for SBSs. Deletions or insertions overlapping with exons taken from the Gff3 file from the reference genome were counted (Kawahara et al., 2013). Only genes disrupted by the breakpoint of inversions or translocations were counted for these two types of variants. Genes in the duplicated regions were counted for each tandem duplication event. We performed gene ontology (GO) analysis on the affected genes using agriGO (http://bioinfo.cau.edu.cn/agriGO/) (Du et al., 2010). In the GO analysis, we used the biological process category.

### Loss-of-Function Mutations

The definition of loss-of-function mutations was adapted from (MacArthur et al., 2012) with minor modifications. We defined loss-of-function mutations as nonsense mutations or SBSs causing changes in the canonical GT/AG intron splicing sites or loss of the start codon, Indels causing frameshifts, and structural variants, including large deletions overlapping genes, and inversions and translocations whose breakpoints fall in genic regions. Tandem duplications were not considered as loss-of-function mutations in this study.

### Heat Maps

To compare the amino acid changes caused by fast-neutron irradiation to those caused by chemical mutagens, such as EMS, we selected one EMS-induced mutant population in rice (Henry et al., 2014) and one ethyl methanesulfonate/ N-ethyl-N-nitrosourea (EMS/ENU)-induced mutant population in *C. elegans* (Thompson et al., 2013), the most comprehensive whole-genome sequenced population of its type in animals. The EMS/ENU-induced *C. elegans* population was created predominantly with either EMS (37% of strains), ENU (13% of strains), or a combination of both (50% of strains) in the published *C. elegans* population (Thompson et al., 2013). We analyzed the nucleotide changes of missense mutations and the resulting amino acid changes of these three FN- or EMS/ENU-induced mutant populations. The analyzed results were incorporated into a matrix format that was used in drawing the heat maps using the R/qplots package (https://www.R-project.org/).

### Cosegregation Assays of the Short Grain Phenotype in Mutant FN1535

A segregating population, including the M_2_ and M_3_ plants derived from FN1535, was used in the cosegregation assay. Fifty plants were used in the assays. Individual M_3_ plants were phenotyped by measuring grain length when seeds were mature. Average seed length was calculated by measuring 10 representative seeds in a row. χ^2^ analyzes were conducted to assay the goodness of fit between the observed the expected values of the segregation ratio. Genomic DNA was isolated from the plants using the CTAB method (see above). Mutation-specific primers Inv/F (5’-ttccgttgctttggaacttt-3’) and Inv/R (5’-cacagcagttttgcacccta-3’) were designed from the flanking sequences of the breakpoint of the inversion on chromosome 5 so that PCR will amplify from the wild-type plant and plants heterozygous at the mutation sites, but not from plants homozygous at the inversion site. Primers targeting the 37 kb deletion region on chromosome 7 are Del/F (5’-catcctcacggctataccaa-3’) and Del/R (5’-ggtgacgacgagcgagag-3’). The actin primers ActF (5’-atccttgtatgctagcggtcga-3’) and ActR (5’-atccaaccggaggatagcatg-3’) were used for DNA quality control. Snapshots of the breakpoints of the inversion on chromosome 5 were taken using Integrative Genomics Viewer (IGV) (Robinson et al., 2011). The diagram of the structure of the mutated gene was modified from the reference genome (Kawahara et al., 2013). PCR was performed with the DreamTaq enzyme (Thermo Scientific).

### KitBase

The open access resource named KitBase (http://kitbase.ucdavis.edu/) integrates genomic data, mutation data, and seed information of the Kitaake rice mutant population. Open source software and tools were used for the development of KitBase. The mutation data of each line were stored in the relational database using MySQL (https://www.mysql.com/). We used the PHP: Hypertext Preprocessor (PHP) scripting language (http://php.net/) to create the web interface and to make the data accessible. Variant Call Format (VCF) files were generated for each type of mutation and embedded in the JBrowse genome browser (Skinner et al., 2009) to visualize the mutations. Standalone BLAST was incorporated into KitBase to facilitate DNA and protein sequence searching (Deng et al., 2007). Both MSU7 LOC gene IDs (http://rice.plantbiology.msu.edu/) and RAP-DB gene IDs (http://rapdb.dna.affrc.go.jp/) were incorporated into KitBase; users can use either when searching KitBase. The seed request webpage facilitates seed distribution. The KitBase server is hosted by the University of California, Davis.

### Accession Numbers

All sequencing data have been deposited to NCBI’s Sequence Read Archive (SRA) (http://www.ncbi.nlm.nih.gov/sra) under accessions listed in Supplemental Data Set 1. Sequencing data are also available from the Joint Genome Institute (JGI) website (http://genome.jgi.doe.gov/). Seed stocks of the Kitaake rice mutant lines of this study are available at KitBase (http://kitbase.ucdavis.edu/kitbase/seed-order).

## Supplemental Data

The following materials are available in the online version of this article.

**Supplemental Figure 1.**
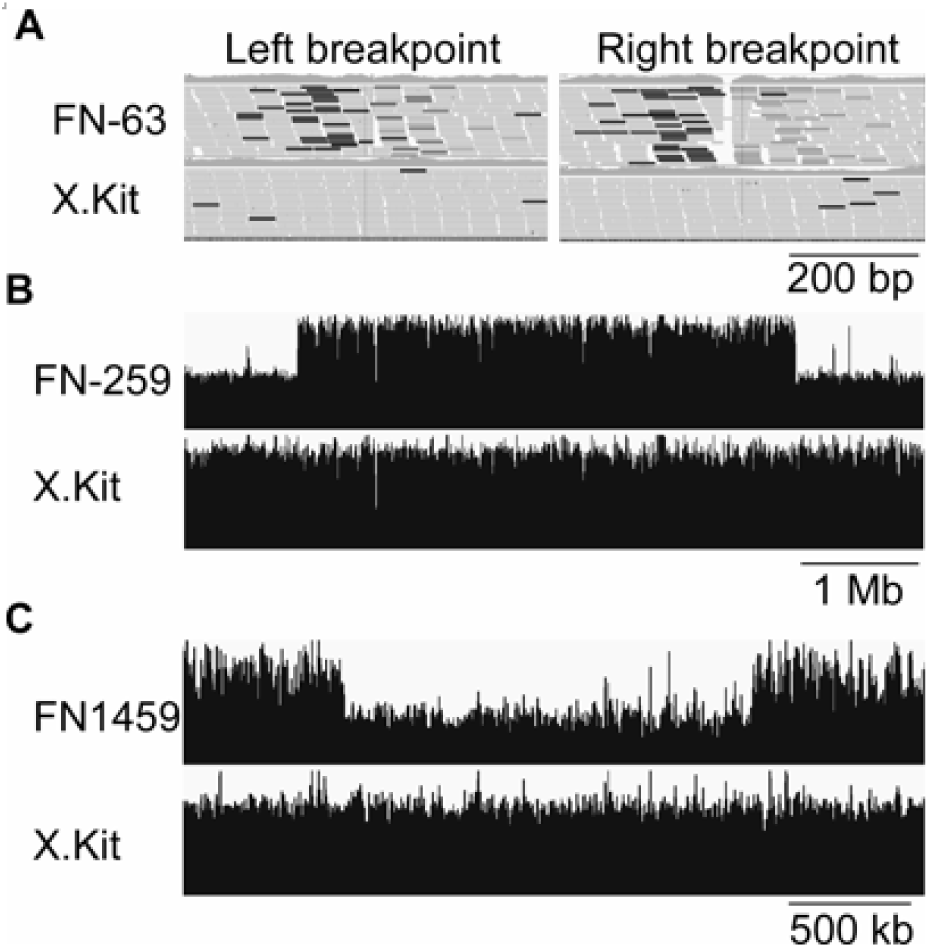
The Largest Inversion, Tandem Duplication, and Deletion Events Detected in the Kitaake Rice Mutant Population. **(A)** The largest inversion in this study was observed in line FN-63. Integrative Genomics Viewer (IGV) screenshots of the two breakpoints of the 36.8 Mb inversion on chromosome one of line FN-63. The dark color indicates the anomalous reads of the inversion. X.Kit is the parental line. **(B)** The largest tandem duplication in this study in line FN-259. An IGV screenshot of the 4.2 Mb tandem duplication on chromosome eight of line FN-259. The black columns indicate the sequence reads. The higher columns in the middle of the top panel indicate the tandem duplication event in line FN-259. X.Kit is the parental line. **(C)** The largest heterozygous deletion in this study in line FN1459. An IGV screenshot of the 1.7 Mb deletion on chromosome five of mutant FN1459. The black columns indicate the sequence reads, and the gap in the middle of the top panel indicates the heterozygous deletion in line FN1459. X.Kit is the parental line.

**Supplemental Figure 2.**
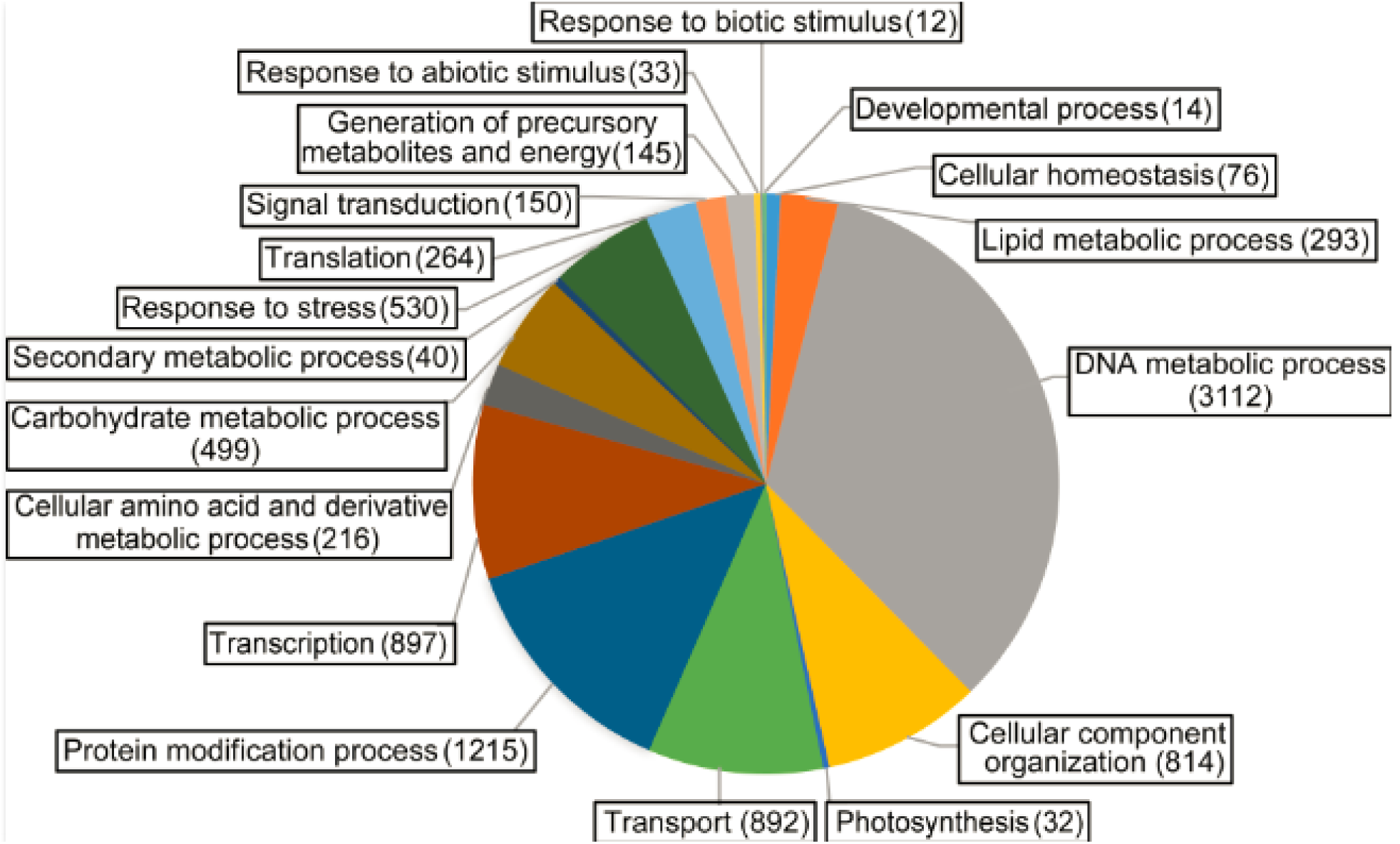
Gene Ontology (GO) Analysis of Affected Genes in the Kitaake Rice Mutant Population. The biological process category was used in the analysis. The number of affected genes assigned to each GO term category is shown in parenthesis.

**Supplemental Figure 3.**
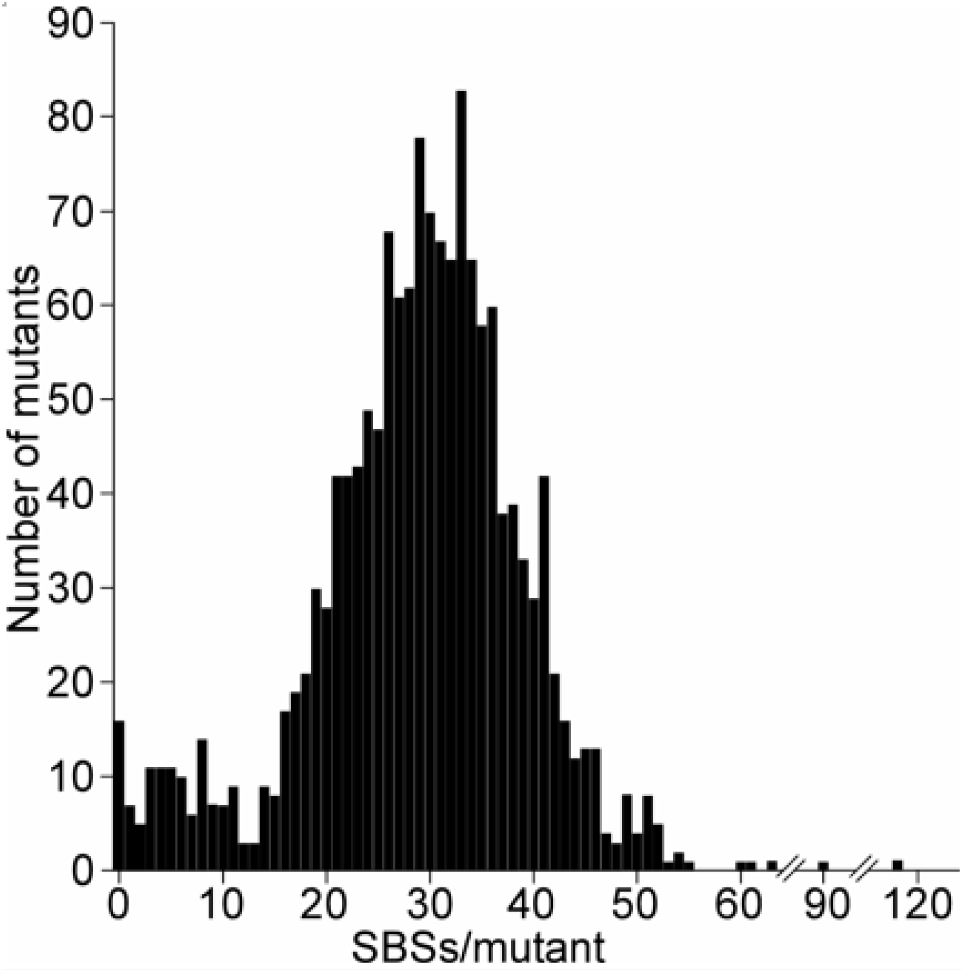
Distribution of the Number of Single Base Substitutions (SBSs) per Line in the Kitaake Rice Mutant Population. The x-axis indicates the number of SBSs per line. The y-axis indicates the number of lines containing that number SBSs.

**Supplemental Figure 4.**
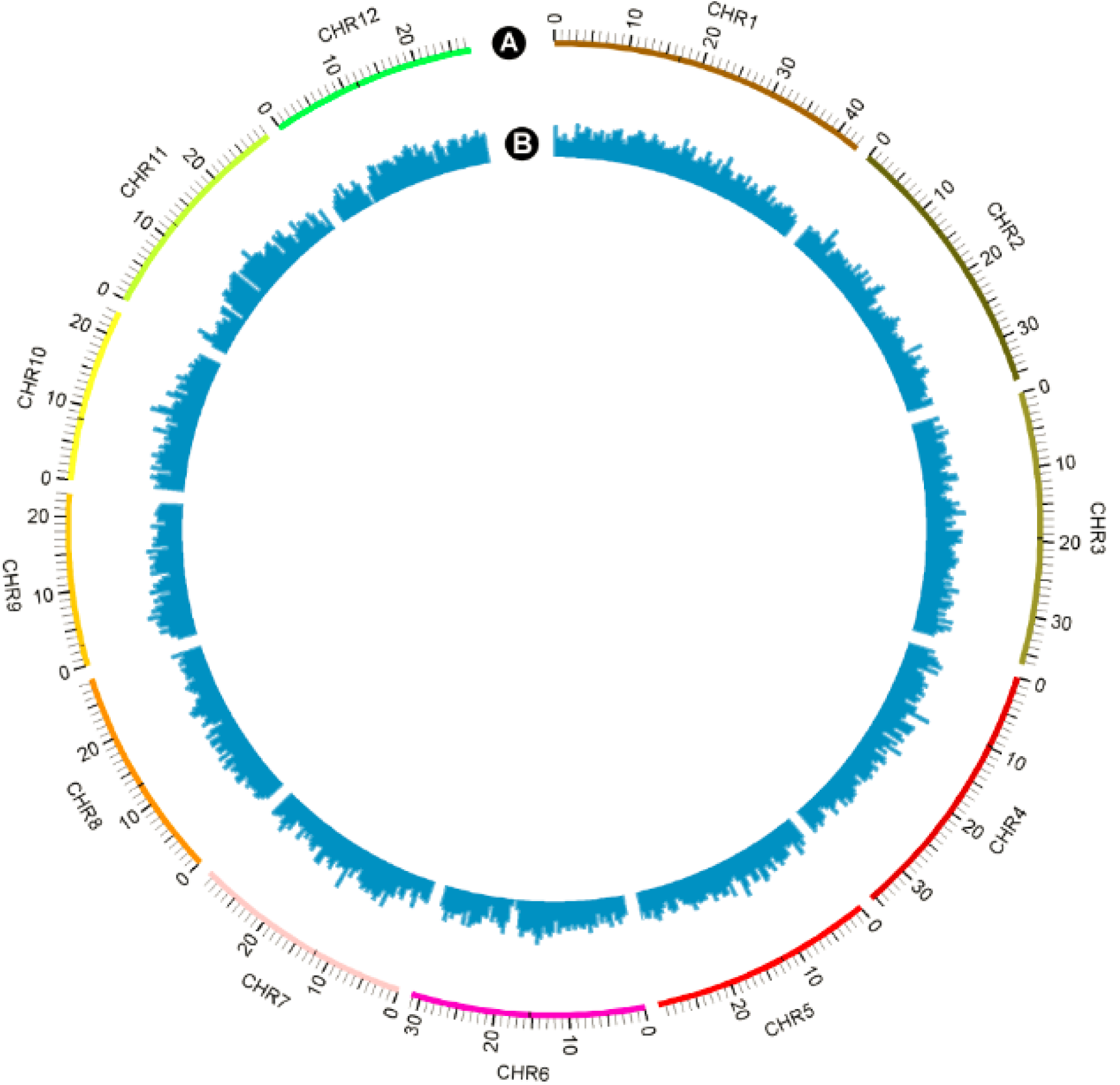
Genome-Wide Distribution of Single Base Substitutions (SBSs) in the Kitaake Rice Mutant Population. **(A)** The twelve rice chromosomes represented on an Mb scale. **(B)** Genome-wide distribution of FN-induced single base substitutions in non-overlapping 500 kb windows. The highest column represents 107 SBSs/500kb.

**Supplemental Figure 5.**
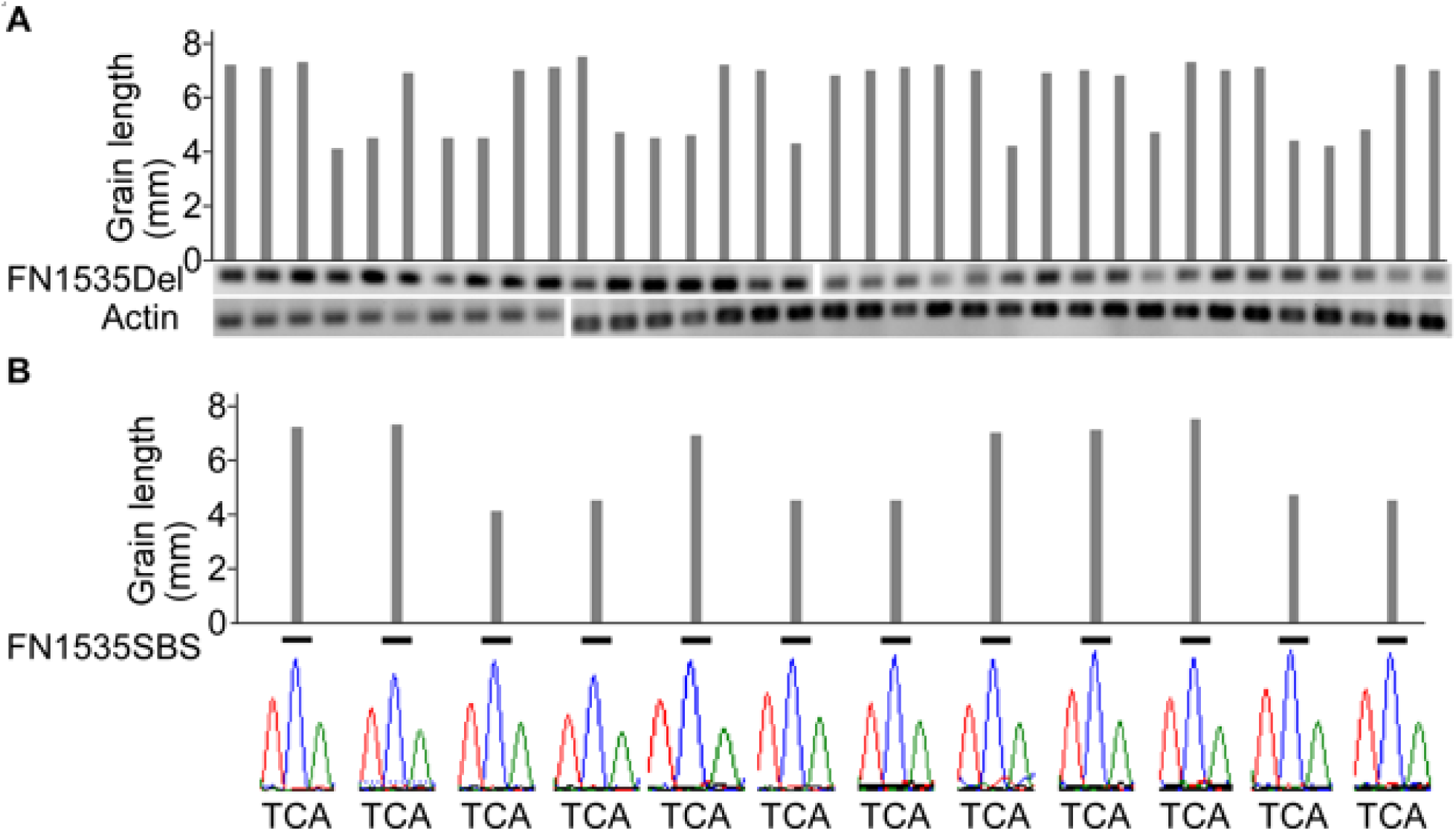
Neither the 37 kb Deletion on Chromosome 7 nor the Single Base Substitution (SBS) on Chromosome 6 of Line FN1535 Cosegregates with the Short-Grain Phenotype. **(A)** The 37 kb deletion on chromosome 7 of line FN1535 does not cosegregate with the short-grain phenotype. Grain length was measured by lining up 10 mature seeds of each plant, and the average grain length was calculated. FN1535Del indicates the PCR results targeting the 37 kb deletion on chromosome 7 of line FN1535. A band indicates that at least one parental allele is in the plant. Actin primers were used for the DNA quality control. The tested population derived from the two M_2_ plants does not have the deletion. **(B)** The single base substitution (SBS) on chromosome 6 of line FN1535 does not cosegregate with the short-grain phenotype. “-” indicates the parental allele. The bottom panel shows the Sanger sequencing results at the position of the SBS in each line.

**Supplemental Table 1.** Translocation Density per Chromosome.

| Chr | Number of Translocations<sup>a</sup> | Translocations/Mb<sup>a</sup> |
| --- | --- | --- |
| Chr1 | 961 | 22.2 |
| Chr2 | 884 | 24.6 |
| Chr3 | 894 | 24.6 |
| Chr4 | 810 | 22.8 |
| Chr5 | 611 | 20.4 |
| Chr6 | 732 | 23.4 |
| Chr7 | 696 | 23.4 |
| Chr8 | 666 | 23.4 |
| Chr9 | 616 | 26.8 |
| Chr10 | 557 | 24.0 |
| Chr11 | 769 | 26.5 |
| Chr12 | 676 | 24.6 |

<sup>a</sup> Both breakpoints of each translocation were used in the calculation.

**Supplemental Table 2.** GO Analysis of Mutated Genes in the Kitaake Rice Mutant Population.

| GO term | Description | Genes | Reference<sup>a</sup> | %<sup>b</sup> |
| --- | --- | --- | --- | --- |
| GO:0005975 | Carbohydrate metabolic process | 499 | 862 | 58 |
| GO:0006091 | Generation of precursor metabolites and energy | 145 | 307 | 47 |
| GO:0006259 | DNA metabolic process | 3,112 | 4,373 | 71 |
| GO:0006350 | Transcription | 897 | 1,691 | 53 |
| GO:0006412 | Translation | 264 | 653 | 40 |
| GO:0006464 | Protein modification process | 1,215 | 1,988 | 61 |
| GO:0006519 | Cellular amino acid & derivative metabolic process | 216 | 406 | 53 |
| GO:0006629 | Lipid metabolic process | 293 | 489 | 60 |
| GO:0006810 | Transport | 892 | 1,627 | 55 |
| GO:0006950 | Response to stress | 530 | 877 | 60 |
| GO:0007165 | Signal transduction | 150 | 251 | 60 |
| GO:0009607 | Response to biotic stimulus | 12 | 18 | 67 |
| GO:0009628 | Response to abiotic stimulus | 33 | 59 | 56 |
| GO:0015979 | Photosynthesis | 32 | 100 | 32 |
| GO:0016043 | Cellular component organization | 814 | 1,215 | 67 |
| GO:0019725 | Cellular homeostasis | 76 | 128 | 59 |
| GO:0019748 | Secondary metabolic process | 40 | 61 | 66 |
| GO:0032502 | Developmental process | 14 | 41 | 34 |

<sup>a</sup> The number of genes belonging to the GO term in the Nipponbare reference genome.
<sup>b</sup> The number of genes in the GO term that are hit in the Kitaake rice mutant population divided by the number of genes in that GO term in Nipponbare reference genome.

**Supplemental Table 3.** Functional Impacts of Single Base Substitutions (SBSs) in the Kitaake Rice Mutant Population.

| Total SBSs | Exons |  |  |  | Untranslated regions | Introns |  | Intergenic |
| --- | --- | --- | --- | --- | --- | --- | --- | --- |
|  | Missense<sup>a</sup> | Nonsense | Readthrough<sup>b</sup> | Synonymous |  | Splicing | Other |  |
| 43,483 | 5,003 | 352 | 55 | 2,127 | 1,368 | 37 | 7,532 | 27,009 |
| Percentage (%) | 11.50 | 0.81 | 0.13 | 4.89 | 3.15 | 0.08 | 17.32 | 62.11 |

<sup>a</sup> Eight start-loss mutations are included.
<sup>b</sup> SBSs that result in new start codons or loss of stop codons.

**Supplemental Table 4.**
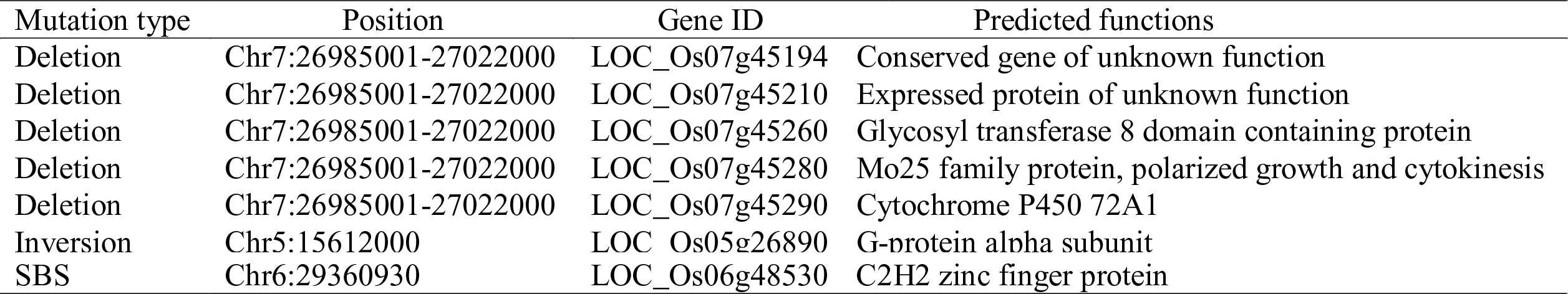
Non-TE Genes Mutated in Line FN1535.

**Supplemental Data Set 1.** Genome Sequencing Summary of Rice Plants Used in This Study.

**Supplemental Data Set 2.** Mutations Identified in the Kitaake Rice Mutant Population.

**Supplemental Data Set 3.** Mutations Selected for Validation.

**Supplemental Data Set 4.** Genes Affected in the Kitaake Rice Mutant Population.

**Supplemental Data Set 5.** Core Eukaryotic Genes Affected in the Kitaake Rice Mutant Population.

**Supplemental Data Set 6.** Genes Mutated by Loss-of-Function Mutations.

**Supplemental Data Set 7.** Genes Mutated by Loss-of-Function Mutations Affecting a Single Gene.

## ACKNOWLEDGMENTS

We thank Patrick E. Canlas, Shuwen Xu, Li Pan, Kira H. Lin, Rick A. Rios, Anton D. Rotter-Sieren, Hans A. Vasquez-Gross, Maria E. Hernandez, Furong Liu, Anna Joe, and Natasha Brown for assistance in genomic DNA isolation and submission, seed organization and data processing, and Drs. Catherine Nelson, Jenny C Mortimer and Brittany Anderton for critical reading of the manuscript. We also thank Drs. Chongyun Fu, Jiandi Xu, and other Ronald lab members for insightful discussions. This work was part of the DOE Joint BioEnergy Institute (http://www.jbei.org) supported by the U. S. Department of Energy, Office of Science, Office of Biological and Environmental Research, through contract DE-AC02-05CH11231 between Lawrence Berkeley National Laboratory and the U. S. Department of Energy. The work conducted by the US Department of Energy Joint Genome Institute (JGI) was supported by the Office of Science of the US Department of Energy under Contract no. DE-AC02-05CH11231. This work was also supported by NIH (GM59962) and NSF (IOS-1237975) to PCR.

## AUTHOR CONTRIBUTIONS

GL, MC, and PR participated in the design of the project, coordination of the project, and data interpretation. GL, RJ and PR drafted and revised the manuscript. MC developed and maintained the mutagenized population. GL, RJ, NP, MC, JM, TW, WS, AL, KJ, JL, PD, RR, DR, DB, YP, KB, and JS performed the sample preparation and sequencing and participated in in-house script development and statistical analyses. All authors read and approved the final manuscript.

## REFERENCES

Abyzov, A., Urban, A.E., Snyder, M., and Gerstein, M. (2011). CNVnator: an approach to discover, genotype, and characterize typical and atypical CNVs from family and population genome sequencing. Genome Res 21, 974-984.

Alonso, J.M., Stepanova, A.N., Leisse, T.J., Kim, C.J., Chen, H.M., Shinn, P., Stevenson, D.K., Zimmerman, J., Barajas, P., Cheuk, R., Gadrinab, C., Heller, C., Jeske, A., Koesema, E., Meyers, C.C., Parker, H., Prednis, L., Ansari, Y., Choy, N., Deen, H., Geralt, M., Hazari, N., Horn, E., Karnes, M., Mulholland, C., Ndubaku, R., Schmidt, I., Guzman, P., Aguilar-Henonin, L., Schmid, M., Weigel, D., Carter, D.E., Marchand, T., Risseeuw, E., Brogden, D., Zeko, A., Crosby, W.L., Berry, C.C., and Ecker, J.R. (2003). Genome-wide Insertional mutagenesis of *Arabidopsis thaliana*. Science 301, 653-657.

Ashburner, M., Ball, C.A., Blake, J.A., Botstein, D., Butler, H., Cherry, J.M., Davis, A.P., Dolinski, K., Dwight, S.S., Eppig, J.T., Harris, M.A., Hill, D.P., Issel-Tarver, L., Kasarskis, A., Lewis, S., Matese, J.C., Richardson, J.E., Ringwald, M., Rubin, G.M., and Sherlock, G. (2000). Gene ontology: tool for the unification of biology. The gene ontology consortium. Nat Genet 25, 25-29.

Ashikari, M., Wu, J., Yano, M., Sasaki, T., and Yoshimura, A. (1999). Rice gibberellin-insensitive dwarf mutant gene Dwarf 1 encodes the alpha-subunit of GTP-binding protein. Proc Natl Acad Sci U S A 96, 10284-10289.

Barampuram, S., and Zhang, Z.J. (2011). Recent advances in plant transformation. Methods Mol Biol 701, 1-35.

Belfield, E.J., Gan, X., Mithani, A., Brown, C., Jiang, C., Franklin, K., Alvey, E., Wibowo, A., Jung, M., Bailey, K., Kalwani, S., Ragoussis, J., Mott, R., and Harberd, N.P. (2012). Genome-wide analysis of mutations in mutant lineages selected following fast-neutron irradiation mutagenesis of *Arabidopsis thaliana*. Genome Res 22, 1306-1315.

Belkadi, A., Bolze, A., Itan, Y., Cobat, A., Vincent, Q.B., Antipenko, A., Shang, L., Boisson, B., Casanova, J.L., and Abel, L. (2015). Whole-genome sequencing is more powerful than whole-exome sequencing for detecting exome variants. Proc Natl Acad Sci U S A 112, 5473-5478.

Biesecker, L.G., Shianna, K.V., and Mullikin, J.C. (2011). Exome sequencing: the expert view. Genome Biol 12, 128.

Bolon, Y.T., Stec, A.O., Michno, J.M., Roessler, J., Bhaskar, P.B., Ries, L., Dobbels, A.A., Campbell, B.W., Young, N.P., Anderson, J.E., Grant, D.M., Orf, J.H., Naeve, S.L., Muehlbauer, G.J., Vance, C.P., and Stupar, R.M. (2014). Genome resilience and prevalence of segmental duplications following fast neutron irradiation of soybean. Genetics 198, 967-981.

Carvalho, C.M., and Lupski, J.R. (2016). Mechanisms underlying structural variant formation in genomic disorders. Nature Rev Genet 17, 224-238.

Chen, K., Wallis, J.W., McLellan, M.D., Larson, D.E., Kalicki, J.M., Pohl, C.S., McGrath, S.D., Wendl, M.C., Zhang, Q., Locke, D.P., Shi, X., Fulton, R.S., Ley, T.J., Wilson, R.K., Ding, L., and Mardis, E.R. (2009). BreakDancer: an algorithm for high-resolution mapping of genomic structural variation. Nat Methods 6, 677-681.

Chen, S., Jin, W., Wang, M., Zhang, F., Zhou, J., Jia, Q., Wu, Y., Liu, F., and Wu, P. (2003). Distribution and characterization of over 1000 T-DNA tags in rice genome. Plant J 36, 105-113.

Cheng, X., Wang, M., Lee, H.-K., Tadege, M., Ratet, P., Udvardi, M., Mysore, K.S., and Wen, J. (2014). An efficient reverse genetics platform in the model legume *Medicago truncatula*. New Phytol 201, 1065-1076.

Chern, M., Xu, Q., Bart, R.S., Bai, W., Ruan, D., Sze-To, W.H., Canlas, P.E., Jain, R., Chen, X., and Ronald, P.C. (2016). A genetic screen identifies a requirement for cysteine-rich-receptor-like kinases in rice NH1 (OsNPR1)-mediated immunity. PLoS Genet 12, e1006049.

Deng, W., Nickle, D.C., Learn, G.H., Maust, B., and Mullins, J.I. (2007). ViroBLAST: a stand-alone BLAST web server for flexible queries of multiple databases and user's datasets. Bioinformatics 23, 2334-2336.

Devos, K.M., and Gale, M.D. (2000). Genome relationships: the grass model in current research. Plant Cell 12, 637-646.

Ding, J., Lu, Q., Ouyang, Y., Mao, H., Zhang, P., Yao, J., Xu, C., Li, X., Xiao, J., and Zhang, Q. (2012). A long noncoding RNA regulates photoperiod-sensitive male sterility, an essential component of hybrid rice. Proc Natl Acad Sci U S A 109, 2654-2659.

Droc, G. (2006). OryGenesDB: a database for rice reverse genetics. Nucleic Acids Res 34, D736-D740.

Du, Z., Zhou, X., Ling, Y., Zhang, Z., and Su, Z. (2010). agriGO: a GO analysis toolkit for the agricultural community. Nucleic Acids Res 38, W64-70.

Goodwin, S., McPherson, J.D., and McCombie, W.R. (2016). Coming of age: ten years of next-generation sequencing technologies. Nature Rev Genet 17, 333-351.

Gross, B.L., and Zhao, Z. (2014). Archaeological and genetic insights into the origins of domesticated rice. Proc Natl Acad Sci U S A 111, 6190-6197.

Henry, I.M., Nagalakshmi, U., Lieberman, M.C., Ngo, K.J., Krasileva, K.V., Vasquez-Gross, H., Akhunova, A., Akhunov, E., Dubcovsky, J., Tai, T.H., and Comai, L. (2014). Efficient genome-wide detection and cataloging of EMS-induced mutations using exome capture and next-generation sequencing. Plant Cell 26, 1382-1397.

Hsing, Y.I., Chern, C.G., Fan, M.J., Lu, P.C., Chen, K.T., Lo, S.F., Sun, P.K., Ho, S.L., Lee, K.W., Wang, Y.C., Huang, W.L., Ko, S.S., Chen, S., Chen, J.L., Chung, C.I., Lin, Y.C., Hour, A.L., Wang, Y.W., Chang, Y.C., Tsai, M.W., Lin, Y.S., Chen, Y.C., Yen, H.M., Li, C.P., Wey, C.K., Tseng, C.S., Lai, M.H., Huang, S.C., Chen, L.J., and Yu, S.M. (2007). A rice gene activation/knockout mutant resource for high throughput functional genomics. Plant Mol Biol 63, 351-364.

Huang, R., Jiang, L., Zheng, J., Wang, T., Wang, H., Huang, Y., and Hong, Z. (2013). Genetic bases of rice grain shape: so many genes, so little known. Trends Plant Sci 18, 218-226.

Huang, X., Kurata, N., Wei, X., Wang, Z.X., Wang, A., Zhao, Q., Zhao, Y., Liu, K., Lu, H., Li, W., Guo, Y., Lu, Y., Zhou, C., Fan, D., Weng, Q., Zhu, C., Huang, T., Zhang, L., Wang, Y., Feng, L., Furuumi, H., Kubo, T., Miyabayashi, T., Yuan, X., Xu, Q., Dong, G., Zhan, Q., Li, C., Fujiyama, A., Toyoda, A., Lu, T., Feng, Q., Qian, Q., Li, J., and Han, B. (2012). A map of rice genome variation reveals the origin of cultivated rice. Nature 490, 497-501.

Itoh, T., Tanaka, T., Barrero, R.A., Yamasaki, C., Fujii, Y., Hilton, P.B., Antonio, B.A., Aono, H., Apweiler, R., Bruskiewich, R., Bureau, T., Burr, F., Costa de Oliveira, A., Fuks, G., Habara, T., Haberer, G., Han, B., Harada, E., Hiraki, A.T., Hirochika, H., Hoen, D., Hokari, H., Hosokawa, S., Hsing, Y.I., Ikawa, H., Ikeo, K., Imanishi, T., Ito, Y., Jaiswal, P., Kanno, M., Kawahara, Y., Kawamura, T., Kawashima, H., Khurana, J.P., Kikuchi, S., Komatsu, S., Koyanagi, K.O., Kubooka, H., Lieberherr, D., Lin, Y.C., Lonsdale, D., Matsumoto, T., Matsuya, A., McCombie, W.R., Messing, J., Miyao, A., Mulder, N., Nagamura, Y., Nam, J., Namiki, N., Numa, H., Nurimoto, S., O'Donovan, C., Ohyanagi, H., Okido, T., Oota, S., Osato, N., Palmer, L.E., Quetier, F., Raghuvanshi, S., Saichi, N., Sakai, H., Sakai, Y., Sakata, K., Sakurai, T., Sato, F., Sato, Y., Schoof, H., Seki, M., Shibata, M., Shimizu, Y., Shinozaki, K., Shinso, Y., Singh, N.K., Smith-White, B., Takeda, J., Tanino, M., Tatusova, T., Thongjuea, S., Todokoro, F., Tsugane, M., Tyagi, A.K., Vanavichit, A., Wang, A., Wing, R.A., Yamaguchi, K., Yamamoto, M., Yamamoto, N., Yu, Y., Zhang, H., Zhao, Q., Higo, K., Burr, B., Gojobori, T., and Sasaki, T. (2007). Curated genome annotation of *Oryza sativa* ssp. *japonica* and comparative genome analysis with *Arabidopsis thaliana*. Genome Res 17, 175-183.

Izawa, T., and Shimamoto, K. (1996). Becoming a model plant: The importance of rice to plant science. Trends Plant Sci 1, 95-99.

Jeon, J.S., Lee, S., Jung, K.H., Jun, S.H., Jeong, D.H., Lee, J., Kim, C., Jang, S., Yang, K., Nam, J., An, K., Han, M.J., Sung, R.J., Choi, H.S., Yu, J.H., Choi, J.H., Cho, S.Y., Cha, S.S., Kim, S.I., and An, G. (2000). T-DNA insertional mutagenesis for functional genomics in rice. Plant J 22, 561-570.

Jiang, W., Zhou, H., Bi, H., Fromm, M., Yang, B., and Weeks, D.P. (2013). Demonstration of CRISPR/Cas9/sgRNA-mediated targeted gene modification in Arabidopsis, tobacco, sorghum and rice. Nucleic Acids Res 41, el88.

Jiao, Y., Burke, J., Chopra, R., Burow, G., Chen, J., Wang, B., Hayes, C., Emendack, Y., Ware, D., and Xin, Z. (2016). A sorghum mutant resource as an efficient platform for gene discovery in grasses. Plant Cell 28, 1551-1562.

Kawahara, Y., de la Bastide, M., Hamilton, J.P., Kanamori, H., McCombie, W.R., Ouyang, S., Schwartz, D.C., Tanaka, T., Wu, J.Z., Zhou, S.G., Childs, K.L., Davidson, R.M., Lin, H.N., Quesada-Ocampo, L., Vaillancourt, B., Sakai, H., Lee, S.S., Kim, J., Numa, H., Itoh, T., Buell, C.R., and Matsumoto, T. (2013). Improvement of the *Oryza sativa* Nipponbare reference genome using next generation sequence and optical map data. Rice 6:4.

Krasileva, K.V., Vasquez-Gross, H.A., Howell, T., Bailey, P., Paraiso, F., Clissold, L., Simmonds, J., Ramirez-Gonzalez, R.H., Wang, X., Borrill, P., Fosker, C., Ayling, S., Phillips, A.L., Uauy, C., and Dubcovsky, J. (2017). Uncovering hidden variation in polyploid wheat. Proc Natl Acad Sci U S A 114, E913-E921.

Krishnan, A., Guiderdoni, E., An, G., Hsing, Y.i.C., Han, C.d., Lee, M.C., Yu, S.M., Upadhyaya, N., Ramachandran, S., Zhang, Q., Sundaresan, V., Hirochika, H., Leung, H., and Pereira, A. (2009). Mutant resources in rice for functional genomics of the grasses. Plant Physiol 149, 165-170.

Krzywinski, M., Schein, J., Birol, I., Connors, J., Gascoyne, R., Horsman, D., Jones, S.J., and Marra, M.A. (2009). Circos: an information aesthetic for comparative genomics. Genome Res 19, 1639-1645.

Lan, Y., Su, N., Shen, Y., Zhang, R., Wu, F., Cheng, Z., Wang, J., Zhang, X., Guo, X., Lei, C., Jiang, L., Mao, L., and Wan, J. (2012). Identification of novel MiRNAs and MiRNA expression profiling during grain development in indica rice. BMC Genomics 13, 264.

Li, C.L., Santhanam, B., Webb, A.N., Zupan, B., and Shaulsky, G. (2016a). Gene discovery by chemical mutagenesis and whole-genome sequencing in *Dictyostelium*. Genome Res 26, 1268-1276.

Li, G., Chern, M., Jain, R., Martin, J.A., Schackwitz, W.S., Jiang, L., Vega-Sanchez, M.E., Lipzen, A.M., Barry, K.W., Schmutz, J., and Ronald, P.C. (2016b). Genome-wide sequencing of 41 rice (*Oryza sativa* L.) mutated lines reveals diverse mutations induced by fast-neutron irradiation. Mol Plant 9, 1078-1081.

Li, H. (2013). Aligning sequence reads, clone sequences and assembly contigs with BWA-MEM. arXiv: 1303.3997.

Li, H., and Durbin, R. (2009). Fast and accurate short read alignment with Burrows-Wheeler transform. Bioinformatics 25, 1754-1760.

Li, T., Liu, B., Spalding, M.H., Weeks, D.P., and Yang, B. (2012). High-efficiency TALEN-based gene editing produces disease-resistant rice. Nature Biotechnol 30, 390-392.

Li, X., Zhang, R., Patena, W., Gang, S.S., Blum, S.R., Ivanova, N., Yue, R., Robertson, J.M., Lefebvre, P.A., Fitz-Gibbon, S.T., Grossman, A.R., and Jonikas, M.C. (2016c). An indexed, mapped mutant library enables reverse genetics studies of biological processes in *Chlamydomonas reinhardtii*. Plant Cell 28, 367-387.

Lowry, D.B., and Willis, J.H. (2010). A widespread chromosomal inversion polymorphism contributes to a major life-history transition, local adaptation, and reproductive isolation. Plos Biol 8, e1000500.

MacArthur, D.G., Balasubramanian, S., Frankish, A., Huang, N., Morris, J., Walter, K., Jostins, L., Habegger, L., Pickrell, J.K., Montgomery, S.B., Albers, C.A., Zhang, Z.D., Conrad, D.F., Lunter, G., Zheng, H., Ayub, Q., DePristo, M.A., Banks, E., Hu, M., Handsaker, R.E., Rosenfeld, J.A., Fromer, M., Jin, M., Mu, X.J., Khurana, E., Ye, K., Kay, M., Saunders, G.I., Suner, M.M., Hunt, T., Barnes, I.H., Amid, C., Carvalho-Silva, D.R., Bignell, A.H., Snow, C., Yngvadottir, B., Bumpstead, S., Cooper, D.N., Xue, Y., Romero, I.G., Wang, J., Li, Y., Gibbs, R.A., McCarroll, S.A., Dermitzakis, E.T., Pritchard, J.K., Barrett, J.C., Harrow, J., Hurles, M.E., Gerstein, M.B., and Tyler-Smith, C. (2012). A systematic survey of loss-of-function variants in human protein-coding genes. Science 335, 823-828.

McCallum, C.M., Comai, L., Greene, E.A., and Henikoff, S. (2000). Targeting induced local lesions IN genomes (TILLING) for plant functional genomics. Plant Physiol 123, 439-442.

McCouch, S.R., Wright, M.H., Tung, C.-W., Maron, L.G., McNally, K.L., Fitzgerald, M., Singh, N., DeClerck, G., Agosto-Perez, F., Korniliev, P., Greenberg, A.J., Naredo, M.E.B., Mercado, S.M.Q., Harrington, S.E., Shi, Y., Branchini, D.A., Kuser-Falcão, P.R., Leung, H., Ebana, K., Yano, M., Eizenga, G., McClung, A., and Mezey, J. (2016). Open access resources for genome-wide association mapping in rice. Nat Commun 7, 10532.

Miao, J., Guo, D., Zhang, J., Huang, Q., Qin, G., Zhang, X., Wan, J., Gu, H., and Qu, L.J. (2013). Targeted mutagenesis in rice using CRISPR-Cas system. Cell Res 23, 1233-1236.

Michael, T.P., and Jackson, S. (2013). The first 50 plant genomes. The Plant Genome 6, 0.

Moscou, M.J., and Bogdanove, A.J. (2009). A simple cipher governs DNA recognition by TAL effectors. Science 326, 1501.

Parra, G., Bradnam, K., Ning, Z., Keane, T., and Korf, I. (2008). Assessing the gene space in draft genomes. Nucleic Acids Res 37, 289-297.

Peng, R., Lin, G., and Li, J. (2016). Potential pitfalls of CRISPR/Cas9-mediated genome editing. FEBS J 283, 1218-1231.

Peters, J.L., Cnudde, F., and Gerats, T. (2003). Forward genetics and map-based cloning approaches. Trends Plant Sci 8, 484-491.

Rausch, T., Zichner, T., Schlattl, A., Stutz, A.M., Benes, V., and Korbel, J.O. (2012). DELLY: structural variant discovery by integrated paired-end and split-read analysis. Bioinformatics 28, i333–i339.

Robinson, J.T., Thorvaldsdottir, H., Winckler, W., Guttman, M., Lander, E.S., Getz, G., and Mesirov, J.P. (2011). Integrative genomics viewer. Nature Biotechnol 29, 24-26.

Sakai, H., Lee, S.S., Tanaka, T., Numa, H., Kim, J., Kawahara, Y., Wakimoto, H., Yang, C.c., Iwamoto, M., Abe, T., Yamada, Y., Muto, A., Inokuchi, H., Ikemura, T., Matsumoto, T., Sasaki, T., and Itoh, T. (2013). Rice annotation project database (RAP-DB): An integrative and interactive database for rice genomics. Plant Cell Physiol 54, e6–e6.

Sallaud, C., Meynard, D., van Boxtel, J., Gay, C., Bes, M., Brizard, J.P., Larmande, P., Ortega, D., Raynal, M., Portefaix, M., Ouwerkerk, P.B., Rueb, S., Delseny, M., and Guiderdoni, E. (2003). Highly efficient production and characterization of T-DNA plants for rice ( Oryza sativa L.) functional genomics. Theor Appl Genet 106, 1396-1408.

Saxena, R.K., Edwards, D., and Varshney, R.K. (2014). Structural variations in plant genomes. Brief Funct Genomics 13, 296-307.

Schneeberger, K. (2014). Using next-generation sequencing to isolate mutant genes from forward genetic screens. Nat Rev Genet 15, 662-676.

Schwessinger, B., Bahar, O., Thomas, N., Holton, N., Nekrasov, V., Ruan, D., Canlas, P.E., Daudi, A., Petzold, C.J., Singan, V.R., Kuo, R., Chovatia, M., Daum, C., Heazlewood, J.L., Zipfel, C., and Ronald, P.C. (2015). Transgenic expression of the dicotyledonous pattern recognition receptor EFR in rice leads to ligand-dependent activation of defense responses. PLoS Pathog 11, e1004809.

Skinner, M.E., Uzilov, A.V., Stein, L.D., Mungall, C.J., and Holmes, I.H. (2009). JBrowse: a next-generation genome browser. Genome Res 19, 1630-1638.

Thompson, O., Edgley, M., Strasbourger, P., Flibotte, S., Ewing, B., Adair, R., Au, V., Chaudhry, I., Fernando, L., Hutter, H., Kieffer, A., Lau, J., Lee, N., Miller, A., Raymant, G., Shen, B., Shendure, J., Taylor, J., Turner, E.H., Hillier, L.W., Moerman, D.G., and Waterston, R.H. (2013). The million mutation project: a new approach to genetics in *Caenorhabditis elegans*. Genome Res 23, 1749-1762.

Ueguchi-Tanaka, M., Fujisawa, Y., Kobayashi, M., Ashikari, M., Iwasaki, Y., Kitano, H., and Matsuoka, M. (2000). Rice dwarf mutant d1, which is defective in the alpha subunit of the heterotrimeric G protein, affects gibberellin signal transduction. Proc Natl Acad Sci U S A 97, 11638-11643.

Wang, L., Zheng, J., Luo, Y., Xu, T., Zhang, Q., Zhang, L., Xu, M., Wan, J., Wang, M.B., Zhang, C., and Fan, Y. (2013a). Construction of a genomewide RNAi mutant library in rice. Plant Biotechnol J 11, 997-1005.

Wang, N.L., Long, T.A., Yao, W., Xiong, L.Z., Zhang, Q.F., and Wu, C.Y. (2013b). Mutant resources for the functional analysis of the rice genome. Mol Plant 6, 596-604.

Wei, F.J., Droc, G., Guiderdoni, E., and Hsing, Y.I.C. (2013). International consortium of rice mutagenesis: resources and beyond. Rice 6, 39.

Weischenfeldt, J., Symmons, O., Spitz, F., and Korbel, J.O. (2013). Phenotypic impact of genomic structural variation: insights from and for human disease. Nature Rev Genet 14, 125-138.

Wu, C., Li, X., Yuan, W., Chen, G., Kilian, A., Li, J., Xu, C., Zhou, D.X., Wang, S., and Zhang, Q. (2003). Development of enhancer trap lines for functional analysis of the rice genome. Plant J 35, 418-427.

Xie, K., Minkenberg, B., and Yang, Y. (2015). Boosting CRISPR/Cas9 multiplex editing capability with the endogenous tRNA-processing system. Proc Natl Acad Sci U S A 112, 3570-3575.

Xu, X., Liu, X., Ge, S., Jensen, J.D., Hu, F.Y., Li, X., Dong, Y., Gutenkunst, R.N., Fang, L., Huang, L., Li, J.X., He, W.M., Zhang, G.J., Zheng, X.M., Zhang, F.M., Li, Y.R., Yu, C., Kristiansen, K., Zhang, X.Q., Wang, J., Wright, M., McCouch, S., Nielsen, R., Wang, J., and Wang, W. (2012). Resequencing 50 accessions of cultivated and wild rice yields markers for identifying agronomically important genes. Nature Biotechnol 30, 105–U157.

Yamamoto, E., Yonemaru, J., Yamamoto, T., and Yano, M. (2012). OGRO: The overview of functionally characterized genes in rice online database. Rice 5, 26.

Yang, S., Wang, L., Huang, J., Zhang, X., Yuan, Y., Chen, J.Q., Hurst, L.D., and Tian, D. (2015). Parent-progeny sequencing indicates higher mutation rates in heterozygotes. Nature 523, 463-467.

Yang, W., Guo, Z., Huang, C., Duan, L., Chen, G., Jiang, N., Fang, W., Feng, H., Xie, W., Lian, X., Wang, G., Luo, Q., Zhang, Q., Liu, Q., and Xiong, L. (2014). Combining high-throughput phenotyping and genome-wide association studies to reveal natural genetic variation in rice. Nat Commun 5, 5087.

Ye, K., Schulz, M.H., Long, Q., Apweiler, R., and Ning, Z. (2009). Pindel: a pattern growth approach to detect break points of large deletions and medium sized insertions from paired-end short reads. Bioinformatics 25, 2865-2871.

Yuan, J.S., Tiller, K.H., Al-Ahmad, H., Stewart, N.R., and Stewart, C.N., Jr. (2008). Plants to power: bioenergy to fuel the future. Trends Plant Sci 13, 421-429.

Zhang, J. (2006). RMD: a rice mutant database for functional analysis of the rice genome. Nucleic Acids Res 34, D745–D748.

Zhang, J., Chen, L.L., Xing, F., Kudrna, D.A., Yao, W., Copetti, D., Mu, T., Li, W., Song, J.M., Xie, W., Lee, S., Talag, J., Shao, L., An, Y., Zhang, C.L., Ouyang, Y., Sun, S., Jiao, W.B., Lv, F., Du, B., Luo, M., Maldonado, C.E., Goicoechea, J.L., Xiong, L., Wu, C., Xing, Y., Zhou, D.X., Yu, S., Zhao, Y., Wang, G., Yu, Y., Luo, Y., Zhou, Z.W., Hurtado, B.E., Danowitz, A., Wing, R.A., and Zhang, Q. (2016). Extensive sequence divergence between the reference genomes of two elite indica rice varieties Zhenshan 97 and Minghui 63. Proc Natl Acad Sci U S A 113, E5163–5171.

Zhang, Z., Mao, L., Chen, H., Bu, F., Li, G., Sun, J., Li, S., Sun, H., Jiao, C., Blakely, R., Pan, J., Cai, R., Luo, R., Van de Peer, Y., Jacobsen, E., Fei, Z., and Huang, S. (2015). Genome-wide mapping of structural variations reveals a copy number variant that determines reproductive morphology in cucumber. Plant Cell 27, 1595-1604.

Zmienko, A., Samelak, A., Kozlowski, P., and Figlerowicz, M. (2014). Copy number polymorphism in plant genomes. Theor Appl Genet 127, 1-18.

